# Consequences of PCA graphs, SNP codings, and PCA variants for elucidating population structure

**DOI:** 10.1101/393611

**Authors:** Hugh G. Gauch, Sheng Qian, Hans-Peter Piepho, Linda Zhou, Rui Chen

## Abstract

SNP datasets are high-dimensional, often with thousands to millions of SNPs and hundreds to thousands of samples or individuals. Accordingly, PCA graphs are frequently used to provide a low-dimensional visualization in order to display and discover patterns in SNP data from humans, animals, plants, and microbes—especially to elucidate population structure. Given the popularity of PCA, one might expect that PCA is understood well and applied effectively. However, our literature survey of 125 representative articles that apply PCA to SNP data shows that three choices have usually been made poorly: PCA graph, SNP coding, and PCA variant. Our main three recommendations are simple and easily implemented: Use PCA biplots, SNP coding 1 for the rare allele and 0 for the common allele, and double-centered PCA (or AMMI1 if main effects are of interest). The ultimate benefit from informed and optimal choices of PCA graph, SNP coding, and PCA variant, is expected to be discovery of more biology, and thereby acceleration of medical, agricultural, and other vital applications.

## Introduction

Single nucleotide polymorphism (SNP) data is common in the genetics and genomics literature, and principal components analysis (PCA) is one of the statistical analyses applied most frequently to SNP data. These PCA analyses serve a multitude of research purposes, including increasing biological understanding, accelerating crop breeding, and improving human medicine. This article focuses on the one research purpose identified in its title, elucidating population structure—although its discussion and citations make evident the broader relevance of the results and principles presented here.

PCA is not a single method that is always done exactly the same way. Rather, three methodological choices are implicated necessarily in each and every PCA analysis and graph of SNP data. They are indicated in this article’s title: the kind of graph produced, the way that SNP reads (A, C, G, or T) are coded numerically, and the transformation applied to the data prior to PCA analysis. These three choices impact which kinds of structure and patterns in SNP data can be displayed and discovered in PCA graphs.

Current practices—as documented by a literature survey of 125 representative articles that apply PCA to SNP data—suffice to justify the well-deserved popularity and abundant success of PCA for elucidating population structure (supporting information S1 Literature Survey). But details matter. Improvements are possible. Greater understanding of the consequences of these three choices opens an opportunity for researchers to make informed and optimal choices, and thereby to gain even more biological insight and practical value from their SNP data. Fortunately, this opportunity comes at a small cost: Changing from one kind of graph to another, or from one SNP coding to another, or from one data transformation to another, as needed in order to optimize PCA analysis, is a simple matter requiring negligible change in procedure, effort, and computation. Effective PCA of SNP data is worthwhile because of numerous vital applications that span microbes, plants, animals, and humans.

The topic of this article is at the interface between statistics and biology. On the one hand, invariant mathematical and statistical principles determine what the choices are for PCA graphs, SNP codings, and PCA variants, as well as the consequences of those choices. On the other hand, variable biological circumstances for individual research projects affect which choices are best. Nevertheless, despite variation from project to project, some choices are best more often than others—even much more often—so some general or default recommendations emerge from this methodological investigation. In order to understand contemporary practices and to identify optimal practices, this article explores three topics: two kinds of PCA graphs, three SNP codings, and six PCA variants.

First, we consider two kinds of PCA graphs. PCA is applicable to a two-way factorial design, that is, a data matrix, and it provides a dual analysis of both the rows and the columns of a matrix. The standard term for a figure showing both is a “biplot.” The contrasting term used here for showing only rows or only columns is a “monoplot.” And our generic term for either a monoplot or a biplot is a “graph.” Biplots were first introduced by Gabriel [1] and have become the norm in countless applications of PCA [2], including ecology and agriculture. Also, biplots are used occasionally for a closely related kind of genomics data, namely gene expression data [3–5]. In the present context, the data matrix has a number of SNPs which have been observed for a number of Individuals, where “Individuals” is our generic term applied to any organisms, such as individual humans, horses, cultivars of wheat, or races of a pathogen. We recommend biplots because they are more informative than monoplots: Visualizing both SNPs *and* Individuals opens new opportunities to display and discover additional interesting biology. Unfortunately, our literature survey did not encounter even a single biplot of SNP data.

Second, we compare three SNP codings. Consider a data matrix comprised of a number of SNPs observed for a number of Individuals. The original reads of nucleotides (A, C, G, and T) constitute categorical data, whereas PCA requires numerical data. But there is no natural and unique method for translating from this categorical data to the required numerical data, so the SNP literature exhibits multiple methods for coding SNP data numerically. Three options for SNP coding are discussed: code the rare allele as 1 and the common allele as 0 for each SNP, the reverse, and a mixture of rare coded 1 or 0 (and hence common 0 or 1). We name them SNP coding rare=1, common=1, and mixed. We recommend SNP coding rare=1 and document its several advantages for elucidating population structure. However, to the best of our knowledge, the consequences of different SNP codings for the appearance and interpretation of PCA graphs has not yet been addressed. Most articles in our survey fail to report which SNP coding was used, and none explicitly specify the recommended SNP coding, which thereby compromises the interpretation and repeatability of published PCA graphs.

Third, we explore six PCA variants. A SNPs-by-Individuals data matrix comprises a two-way factorial design. Although analysis of variance (ANOVA) has not been used in the present context of PCA analysis of SNP data, it provides important insight by distinguishing three sources of variation that have quite different biological meanings: the SNP main effects, Individual main effects, and SNP-by-Individual (S×I) interaction effects [6]. The six PCA variants discussed here emerge from the application of various data transformations prior to PCA analysis. The main three variants result from subtracting SNP or Individual or both effects from the data matrix prior to PCA, and that subtraction is called centering. Hence, these variants are called SNP-Centered, Individual-Centered, and Double-Centered PCA (DC-PCA). Three additional PCA variants are also mentioned briefly: SNP-Standardized, Individual-Standardized, and Grand-Mean-Centered PCA. We recommend DC-PCA for several reasons, including that DC-PCA uniquely has a single and simple set of interpretive principles, and that graphs from other PCA variants can look wildly different from DC-PCA. However, most articles in our survey fail to report which PCA variant was used, although they often report which software was used, so again that compromises the interpretation and repeatability of published PCA results.

Our literature survey encountered no clear implementation of *even one* of our three recommendations—biplots, SNP coding rare=1, and PCA variant DC-PCA. Consequently, the likelihood that any published PCA analysis of SNP data has yet implemented *all three* recommendations is quite small. Awareness of the consequences of these three choices—which are made in *every* PCA analysis of SNP data necessarily—creates new opportunities to elucidate population structure more effectively.

## Results

### Choices between two PCA graphs

The increase in biological information that is achieved upon upgrading from a monoplot to the recommended biplot can be illustrated by discussing Fig 1 twice: first as a monoplot by considering only its left half, and then as a biplot by considering the whole figure. In this first subsection of the results, we merely state *that* we chose SNP coding rare=1 and PCA variant DC-PCA, but the following two subsections will explain the exact meanings of these choices and the reasons *why* we prefer them. To distinguish DC-PCA from other variants of PCA, its principal components are called interaction principle components (IPCs). By definition, after row and column averages have been subtracted from a data matrix, what remains is the matrix of interactions, as will be explained in greater detail in the third subsection and the appendix.

In its *left half*, Fig 1 shows a typical PCA graph seen in the literature on population structure, with different groups or subsets indicated by different colors, namely oats (*Avena sativa* L.) in this case. Kathy Esvelt Klos kindly shared with us this dataset with 635 oat lines by 1341 SNPs (personal correspondence, 4 June 2018). All SNPs are biallelic, and there are no missing data. Experienced oat breeders had classified the 635 oats into three groups [7]. The 411 spring oats are shown here in green, 103 world diversity oats in blue, and 121 winter oats, which are also called Southern US oats, in red. The spring (green) and winter (red) oats are expected to cluster and to contrast, whereas the world diversity oats (blue) are heterogeneous and are expected to be less clustered. Indeed, this figure visualizes that expected population structure, with IPC1 concentrating spring oats at the left and winter oats at the right. Supporting information S2 Oat Mixed contains this dataset after shifting its values to 0 and 1, and S3 Oat Rare1 has the version after also reversing SNP polarity as needed to code the rare allele as 1.

**Fig 1.**
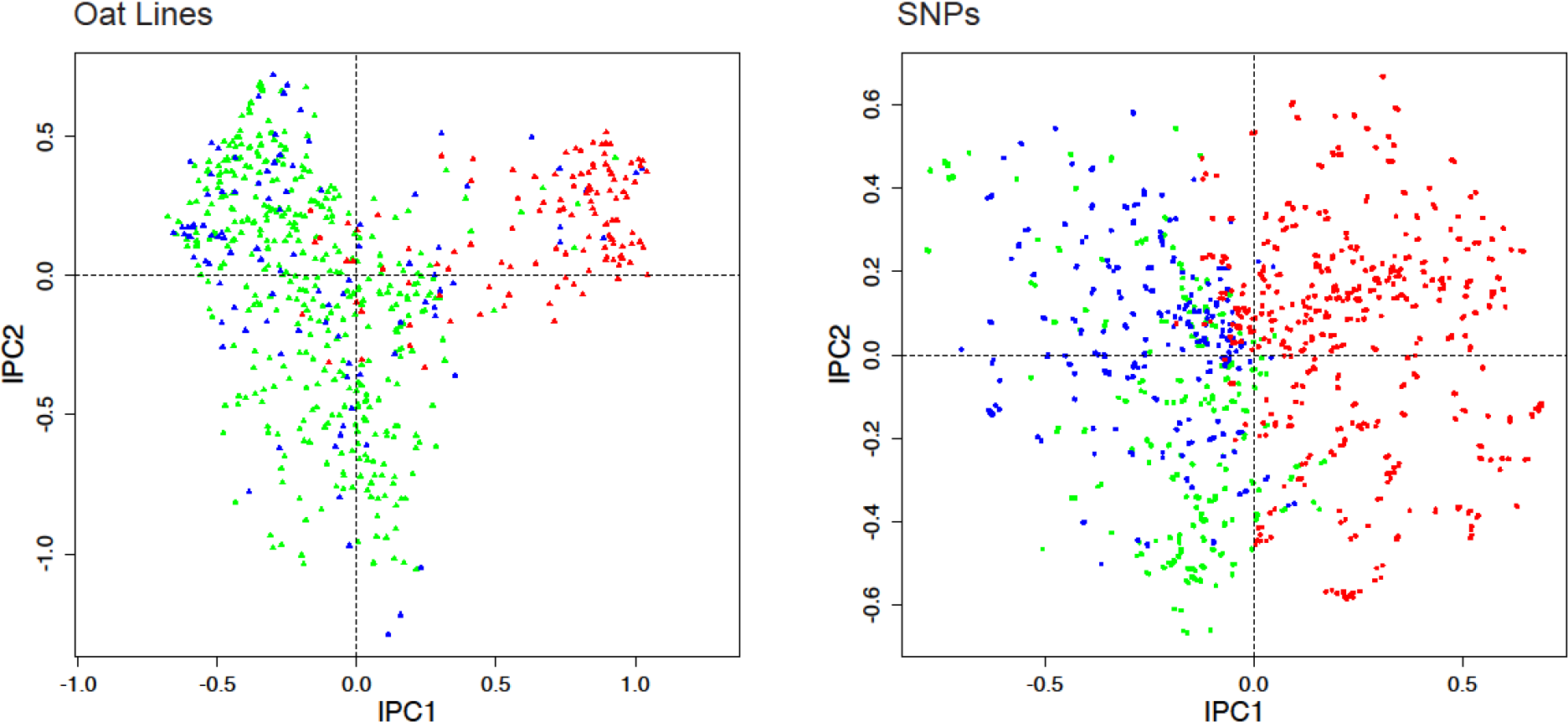
DC-PCA biplot for the oat data, using SNP coding rare=1 and expert knowledge of the oats. To reduce clutter the biplot uses two panels, with oat lines on the left and SNPs on the right. The 635 oat lines are classified in three groups: 411 spring oats shown in green, 103 world diversity oats in blue, and 121 winter oats in red. Likewise, the 1341 SNPs are classified in three groups based on which oat group has the highest percentage of the rare allele: 372 highest in spring oats shown in green, 345 highest in world diversity oats in blue, and 624 highest in winter oats in red.

In the *entire figure*, Fig 1 shows the biplot for this oat dataset, which includes results not only for Individuals, but also for SNPs. This biplot presents Individuals and SNPs in two adjacent panels in order to reduce clutter (whereas biplots with fewer points typically distinguish the two kinds of points with different colors or shapes, and combine both in a single graph, as illustrated in the following subsection). In order for biplots to be useful in the present context of elucidating population structure, methods must be found to color the points in both panels with a coherent color scheme that has a unified biological meaning. Then the joint structure of SNPs-and-Individuals, which is biologically highly relevant, becomes evident. Of course, since biplots have not yet been used in the literature on PCA analyses of SNP data, this literature does not present any methods to upgrade from a monoplot to a biplot with a coherent color scheme in both panels. Fortunately, precedents in the ecological literature, which are directly applicable to the genomics literature, show how to upgrade by two fundamentally different methods that are deeply complementary and synergistic: One method utilizes expert knowledge, and the other automated statistics.

Beginning with the approach using expert knowledge, we devised a method that transfers expert knowledge of the oats to the SNPs in order to classify the 1341 SNPs into three corresponding groups, again using the same color scheme that had been applied to the 635 oats. It is trivially simple:The color assignment for each SNP is based on which of the three oat groups has the highest percentage of the rare allele, and the outcomes were: 372 SNPs colored green had the highest percentage for spring oats, 345 colored blue were highest for world diversity oats, and 624 colored red were highest for winter oats. Thereby, this biplot shows what a monoplot cannot possibly show, the joint structure of oats-and-SNPs, with green mostly on the left and red mostly on the right *in both panels*. Furthermore, when some SNPs have been associated with traits of agricultural or medical importance, such as the 25 SNP associations with agriculturally important traits found in this oat dataset [7], those SNPs can be identified to make the biplot more biologically informative.

Progressing to the approach using automated statistics, we must begin by discussing the PCA arch. When interpreting a PCA graph, such as the biplot for oats in Fig 1, the PCA arch distortion, also called the horseshoe effect, complicates the interpretation of PCA graphs because an underlying one-dimensional gradient (from spring to winter oats in this case) is distorted into an arch in the PC1-PC2 plane. Consequently, awareness of this distortion is necessary for proper interpretation of PCA biplots and graphs. The arch distortion has been well known for decades by archaeologists (D. G. Kendall pages 215–252 in [8]) and ecologists [9, 10]. David Morrison discussed the arch in the context of genomics data [11]; also see [12]. However, his extensive search of the genomics literature found only two papers that discuss the PCA arch distortion [13, 14]. Those papers and his blogposts have not yet succeeded in making this distortion well known in genomics (personal correspondence, David Morrison, 18 February 2018).

Morrison’s initial example is a simple matrix with 1s along the diagonal and 0s elsewhere to represent a SNP dataset that is structured by a single environmental or causal gradient. He showed that its PCA graph has the typical arch distortion. Thereby, Morrison demonstrated a critical implication: *A diagonal data structure implies a PCA arch.* Importantly, the reverse implication also holds, as the ecological literature makes clear, and it merits greater attention in the genomics literature: *A PCA arch implies a diagonal data structure.* To understand this arch distortion, Morrison drew upon the ecological literature, which has a long history of analysis of that arch, unlike the genomics literature. Accordingly, the following one-paragraph review of the pertinent ecology is relevant and helpful in the present context of SNP data.

Ecologists have repeatedly found along an environmental gradient—such as low to high altitude, or dry to wet conditions—a turnover in species abundances, with each species having its own particular environmental preference (Figs 1.3, 3.2 to 3.10, and 6.3 in [9]), and this data structure causes PCA to have an arch distortion (Fig 4.7 in [9]). If the species are listed in an ecologically irrelevant manner, such as alphabetically, and the samples are also listed arbitrarily, such as the order in which they happened to have been collected, the resulting species-by-samples data matrix lacks any discernable structure (Table 1.2 in [9]). Ecologists have developed two sorts of procedures for rearranging the order of matrix rows and columns in order to make structure obvious, using either expert knowledge or automated statistics. First, if the samples are arranged according to knowledge of their environmental conditions (say from dry to wet), and likewise the species are arranged according to their known environmental preferences, then the resulting species-by-samples data matrix has a diagonal structure, with its largest values concentrated along the matrix diagonal (Table 1.3 in [9]). Even if expert knowledge is available for only one matrix dimension, such as the species environmental preferences, simple methods can obtain a corresponding ordering of the samples (such as weighted averages, Table 4.4 in [9]), and thereby obtain an arranged matrix with diagonal structure. Second, provided that a dataset has been structured by a single dominant environmental or causal factor—as suggested by the presence of a PCA arch—even if that factor is not known, the diagonal structure can still be discovered and displayed by automated statistics (Table 1.4 as contrasted with Table 1.2 in [9]). This analysis orders matrix rows and columns by their ranked scores for the first component of correspondence analysis (CA, also called “reciprocal averaging” among ecologists, [15, 16]), which is related to PCA and also involves singular value decomposition (SVD) but uses chi-squared distances rather than Euclidean distances. Incidentally, PCA cannot be substituted for CA to get an arranged matrix because it concentrates small values in the middle of the matrix and thereby fails to display diagonal structure (Fig 4.9 in [9]). Importantly, diagonal structure in a data matrix can be discerned and displayed using either expert ecological knowledge or automated statistical analysis, and ordinarily these two approaches closely agree (Tables 1.3 and 1.4 in [9]). Consequently, three things go together, necessarily and inseparably: a major causal factor or gradient that imposes joint structure on the rows and columns of a data matrix, a data matrix that can be arranged to concentrate large values along its diagonal, and a PCA arch. This is one story told three ways; given any one of these features, all three will occur. Although this unified story has been familiar to ecologists for decades, its relevance for SNP research has not yet been noticed.

The oat lines in Fig 1 show an arch, or actually an upside-down arch in this case. This arch happens to be a filled arch, with many points inside the arch; but PCA arches can also be clear, with few or no points inside (such as Fig 4 in [17]). Our literature survey found that about 80% of PCA graphs of SNP data have an evident arch. For several reasons that emerge in the remainder of this article, PCA arches are a potent source of both problems and opportunities.

The foremost opportunity is a novel method to give both panels of a biplot a coherent color scheme by means of automated statistics, in marked contrast to the above method based on expert knowledge. Because Fig 1 for the oat dataset has the PCA arch distortion, one may rightly expect that this dataset has a diagonal structure that becomes evident after its matrix rows and columns are arranged in rank order of their CA1 scores. Fig 2 shows that such is the case. Data values of 1 for the rare allele are shown in dark blue, and values of 0 for the common allele in light blue. Note that the upper left and lower right corners of this matrix are decidedly darker than the other two corners: SNPs at the left have a concentration of the rare allele (dark blue) for spring oats and a concentration of the common allele (light blue) for winter oats; the reverse also holds for SNPs at the right. The bottom fifth of this arranged matrix contains mostly the 121 oat lines classified as winter oats, which clearly differ in their SNP data from the top four-fifths of this matrix that contains mostly the 411 spring oats and 103 diversity oats. A simple proof that the joint structure of oats-and-SNPs in Fig 2 reflects real structure in the data, rather than arises as an artifact of the CA1 ordering, is that randomizing the order of the oat lines for each SNP individually and then repeating the CA1 ordering makes the matrix structure completely disappear. Of course, the data matrix as received, with SNPs and oats listed in an ecologically arbitrary order, shows no diagonal structure. The diagonal structure in Fig 2 is modest because of the filled PCA arch in Fig 1, whereas diagonal structure is more striking for other datasets with a clear PCA arch. Our literature survey found that of the 80% of PCA graphs of SNP data that have an evident arch, about half are filled and half are clear. This oat example with a heavily filled arch demonstrates that CA1 ordering works even for rather challenging datasets.

**Fig 2.**
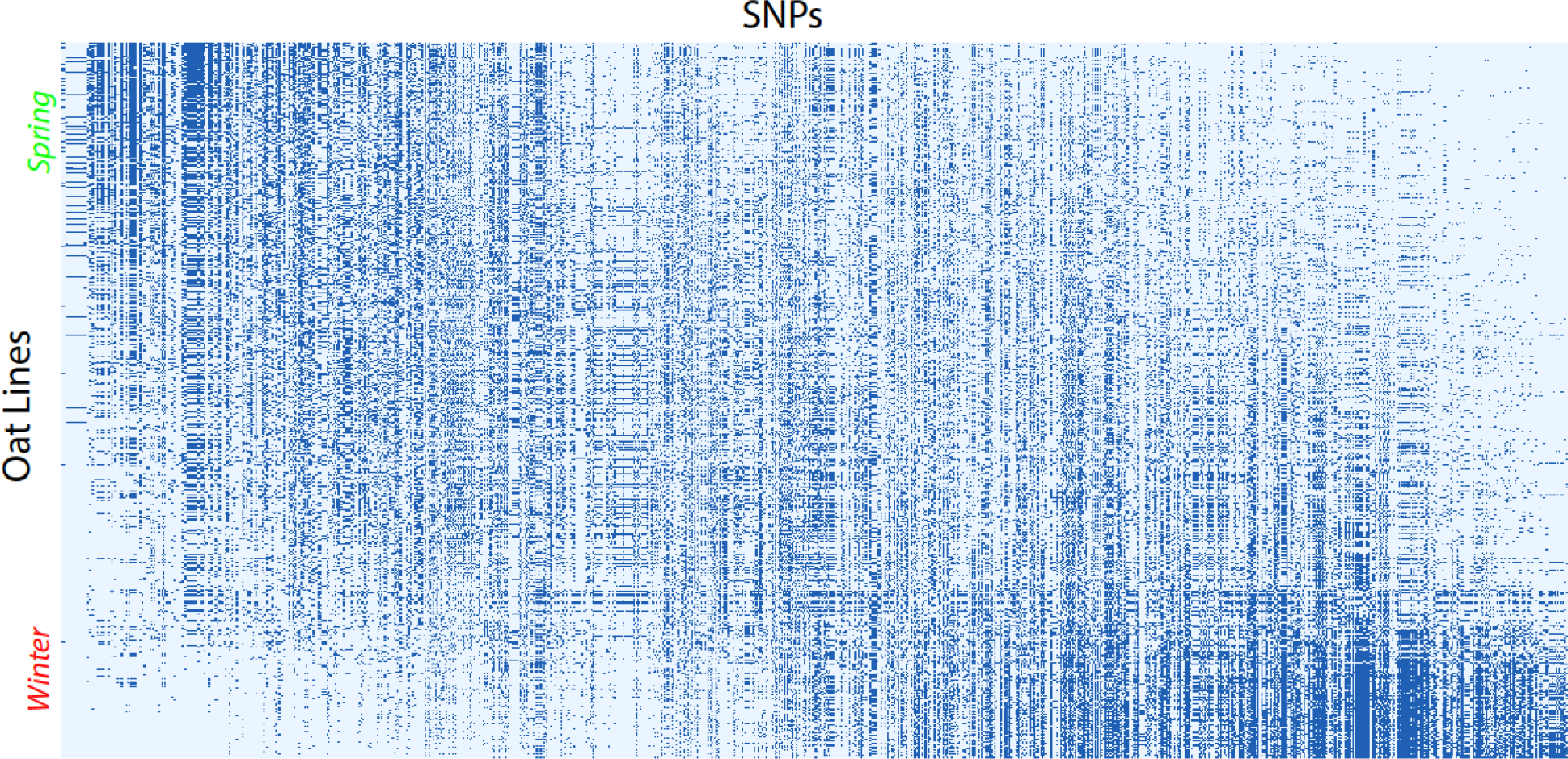
CA1 arranged matrix for 635 oat lines in rows and 1341 SNPs in columns. Spring oats are concentrated at the top of this matrix, and winter oats at the bottom. Correspondingly, SNPs at the left have high percentages of the rare allele in spring oats, and SNPs at the right have high percentages of the rare allele in winter oats.

Furthermore, the PCA arch in Fig 1 implies the diagonal matrix structure in Fig 2, which in turn implies a guaranteed opportunity to discover and display the joint structure of SNPs-and-Individuals by a biplot, even without any expert knowledge of either the SNPs or the Individuals. Indeed, this works. Fig 3 shows exactly the same biplot as Fig 1, except that the Individuals and SNPs have been colored by automated statistics, rather than expert knowledge. This automated method is based on the same ranked CA1 scores for oat lines and for SNPs that were also used to construct Fig 2. The 635 oat lines are subdivided into 5 equal groups of 127 lines according to CA1 order from top to bottom (spring to winter oats) in Fig 2, and likewise the 1341 SNPs form 5 groups of 268 SNPs (plus 1 extra for the last group) from left to right. The corresponding color scheme is: dark green, light green, black, pink, and red. For example, dark green triangles in the left panel are oat lines 1–127, and dark green dots in the right panel are SNPs 1–268. Of course, we could have chosen 3 or 7 equal groups instead of 5 for Fig 3, but we found that 5 provided some extra resolution while still maintaining the color scheme of green (light or dark) for spring oats and red (or pink) for winter oats that was used in Fig 1.

**Fig 3.**
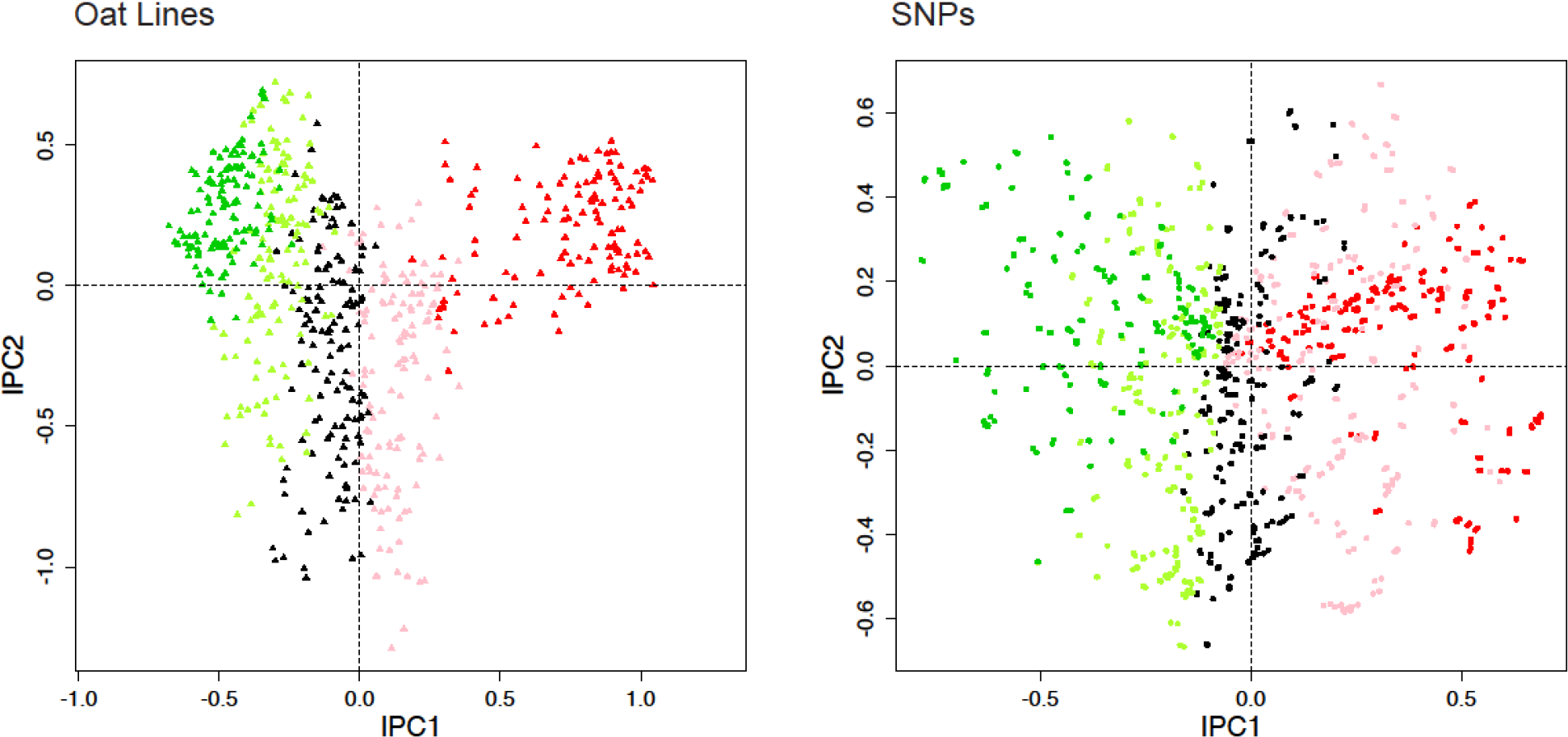
DC-PCA biplot for the oat data, using SNP coding rare=1 and automated statistical analysis providing CA1 order. On the left, the 635 oat lines are subdivided into 5 equal groups according to CA1 order from top to bottom (spring to winter oats) in Fig 2, and these groups are colored dark green, light green, black, pink, and red. On the right, the same is done for the 1341 SNPs from left to right in Fig 2.

The gradient in Fig 3 from Individuals marked dark green at the left to those marked red at the right clearly corresponds to the gradient in Fig 1 from Individuals marked green at the left to those marked red at the right, and the same applies to the SNPs. Therefore, both expert knowledge and automated statistics find the same joint structure of oats-and-SNPs in this dataset. The cause of this joint structure is the climatic and agroecological contrast between southern and northern US locations, which calls for plant breeders to provide winter oats (also called Southern US oats) in the south and spring oats in the north, and that agroecological contrast also corresponds to genome-wide differences in SNPs, as Figs 2 and 3 demonstrate.

Occasional discrepancies may occur amidst an overall similarity when comparing the population structure shown by expert knowledge and by automated statistics, as in Figs 1 and 3, and such discrepancies may be of biological interest. For instance, in the left panel of Fig 1 based on oat breeders’ expert knowledge, consider the anomalous rightmost green point for oat line 128 (CI8000-4) classified as a spring oat but surrounded by mostly red points, or the anomalous leftmost red point for oat line 577 (UPFA_22_Temprana) classified as a winter oat but surrounded by mostly green points—and for present purposes ignore the blue points for world diversity oats that are not expected to cluster. For comparison, Fig 3 based on automated statistics, completely apart from any expert knowledge, avoids these anomalies, with oat 128 colored red like its neighbors, and oat 577 colored green like its neighbors. Furthermore, Fig 2 shows that oat 128, which is near the bottom of this CA1 arranged matrix, has the genome-wide SNP profile characteristic of winter oats, namely light blue at the left and dark blue at the right, meaning the common allele coded 0 for SNPs at the left and the rare allele coded 1 for SNPs at the right; whereas oat 577 near the top has the opposite genome-wide SNP profile characteristic of spring oats. Consequently, the discrepancies between Figs 1 and 3 might prompt oat breeders to review their background information on these anomalous oat lines, and to consider whether oat 128 should be re-classified as a winter oat, and oat 577 as a spring oat. Additional anomalous oats in Fig 1 might also merit reevaluation. The genome-wide information on SNPs in Fig 2 and the right panel of Figs 1 and 3 is useful not only to appreciate the big picture of SNP structure in the entire population, but also to detect anomalies and focus on specific oat lines of particular interest. Thereby, an automated, data-driven approach may help researchers to cross check and refine expert knowledge, thereby making the final results more objective, reliable, and confident.

In review, it is axiomatic that a biplot is more informative than a monoplot. A biplot can, but a monoplot cannot, display and discover joint structure in the Individuals-and-SNPs. Either expert biological knowledge or automated statistical analysis can be used to reveal patterns in PCA biplots—provided that the data are structured by a single dominant causal factor, or equivalently that the PCA graph has the arch distortion. Expert knowledge and automated statistics are deeply complementary, so this combination makes results more reliable and objective. Unfortunately, our literature survey of PCA analyses of SNP data encountered no biplots. However, the simple change of switching from monoplots of Individuals only to biplots of both Individuals and SNPs offers new opportunities to display joint structure, which can increase biological insight. The following section on Materials and Methods provides additional information on the construction and interpretation of PCA graphs.

### Choices between three SNP codings

The biological information displayed in a biplot using the recommended SNP coding rare=1 is clear because this coding uniquely orients the information in both panels of a biplot in the same way, thereby facilitating straightforward and intuitive interpretation—especially of the joint structure of SNPs-and-Individuals. Given the recommendation of biplots in the previous subsection, this subsection evaluates SNP codings primarily in terms of their consequences for the appearance, interpretation, and utility of biplots. However, to accommodate researchers who may prefer monoplots, at least on some occasions, we also discuss the consequences of SNP codings for monoplots. Also, given the recommendation of PCA variant DC-PCA in the next subsection, this subsection explores the consequences of SNP codings for only DC-PCA, leaving further exploration to the next subsection.

One option, here called SNP coding rare=1, is to code the rare allele as 1 and the common allele as 0. This coding is of special interest for reasons that emerge momentarily. Another option, here called SNP coding common=1, is the opposite:to code the common allele as 1 and the rare allele as 0. This coding is the default in TASSEL, which is widely used for crop plants ([18]; Peter Bradbury personal correspondence, 23 April 2018). The third and final possibility, here called SNP coding mixed, is to code the alleles in some other manner that yields a mixture of rare and common alleles coded as 1 (and likewise as 0). For example, the variant call format (VCF) distinguishes one Individual as the reference genome, and then for each SNP it assigns 0 to the allele of the reference genome and 1 to the other non-reference allele. VCF is popular because it was developed for the 1000 Genomes Project in human genetics [19], and subsequently has been adopted widely.

The SNP coding rare=1 can be generalized for SNP datasets having more than the 2 codes of biallelic data. If need be, express the data as a set of consecutive integer values starting with 0, such as 0, 1, 2, and 3. Then recode each SNP to give the rarest allele the highest value, and so on, and lastly give the most common allele the lowest value of 0. Similarly, for a diploid species with three codes—one for each of the two homozygotes and another for the heterozygote—if need be, express the data as 0 and 2 for the homozygotes and 1 for the heterozygote. Then recode each SNP to give the rarest homozygote the value 2 and the most common homozygote the value 0. These ideas can be elaborated for polyploids such as hexaploid (AABBDD) bread wheat, *Triticum aestivum* (L.). The other two codings can also be generalized.

We begin our exploration of the consequences of SNP coding with the simple dataset that Morrison used as the initial example in his first blogpost on the PCA arch distortion in genomics data [11]. His toy dataset has 20 Individuals in its rows and 24 SNPs in its columns. Fig 4 shows three versions of that dataset, with zeroes denoted by dots in order to make the ones readily visible. SNP coding rare=1 is shown on the left, which is Morrison’s original example. SNP coding common=1 is shown in the middle, which is the reverse of the original version. And SNP coding mixed is on the right, which alternates rare=1 and common=1. In all three matrices, red is used for SNPs or columns using rare=1, and green for common=1. The convention adopted here is to number matrix columns from left to right, and number matrix rows from top to bottom, starting with 1 for the first column and the first row.

**Fig 4.**
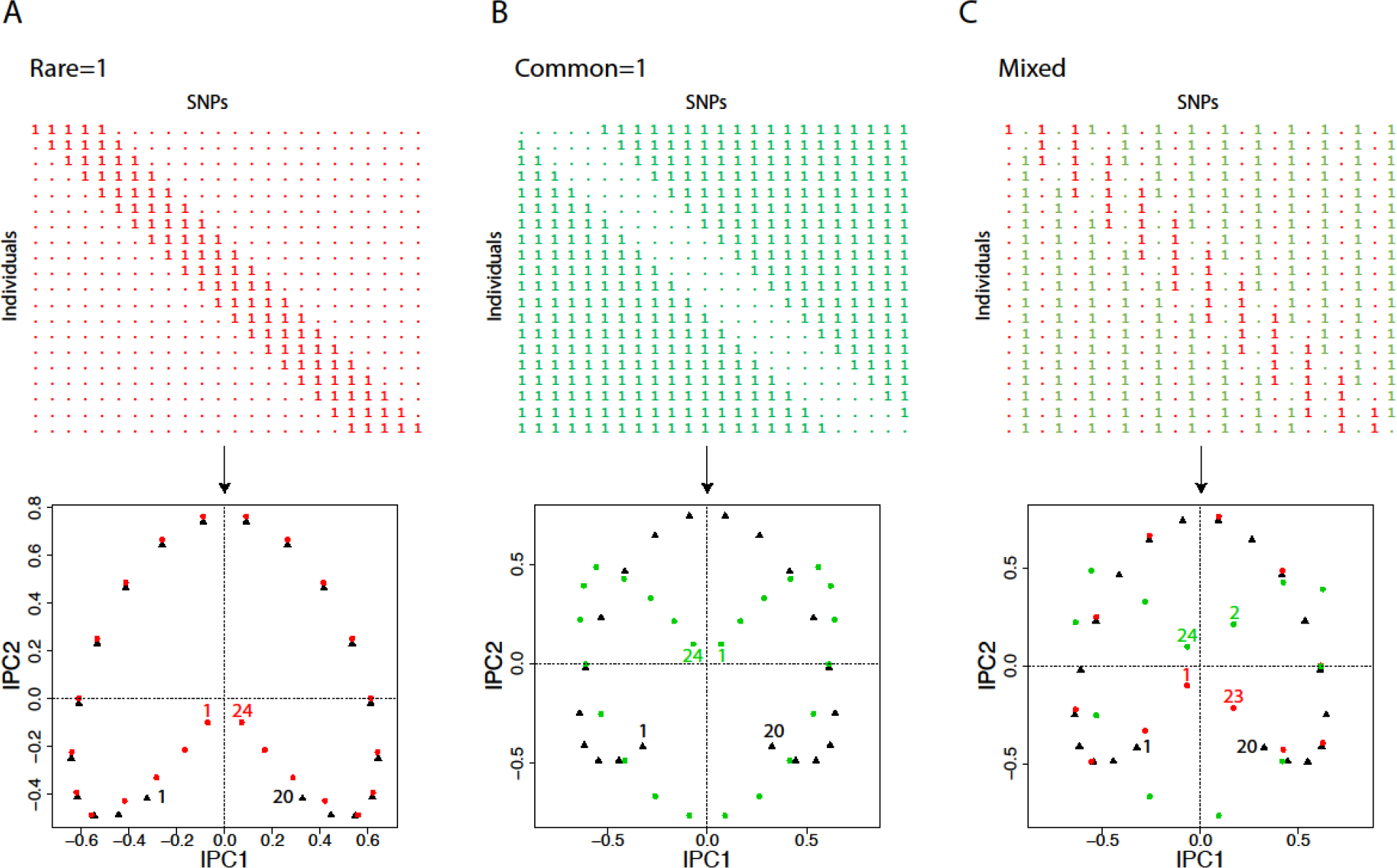
A simple matrix and its DC-PCA biplot, using three SNP codings. The matrix has 20 Individuals in its rows and 24 SNPs in its columns, with SNPs (columns) coded either red for SNP coding rare=1, or green for common=1. The biplots show the 20 Individuals as black triangles, and the 24 SNPs as dots colored either red for rare=1, or green for common=1. Selected points in the biplots are identified: Individuals are numbered 1 to 20 from top to bottom in the data matrix, and SNPs 1 to 24 from left to right. (A) For SNP coding rare=1, despite the arch distortion in the biplot, related Individuals and SNPs track each other in a clear and intuitive manner. (B) For SNP coding common=1, the arch for SNPs is rotated 180° relative to the arch for Individuals, which is confusing. (C) For SNP coding mixed, those SNPs with coding rare=1 (red dots) track the Individuals (black triangles), whereas those SNPs with coding common=1 (green dots) are rotated 180° relative to the other two arches. Taken together without distinguishing red from green, the dots for the SNPs approximate a circle rather than an arch, so awareness of the arch distortion would not suffice to interpret this very confusing biplot properly.

DC-PCA biplots for these three datasets are also shown in this figure. On the left, the PCA arch for the 20 individuals shown with black triangles was included in Morrison’s first blogpost. But this figure presents a biplot, so it also includes the PCA arch for the 24 SNPs shown with red dots. The concentration of ones along the matrix diagonal constitutes a single gradient with evident joint structure that involves *both* Individuals and SNPs. However, DC-PCA has distorted that single gradient into an arch with its ends involuted toward the middle for Individuals 1 to 20, and likewise for SNPs 1 to 24. This arch is a problem that complicates the interpretation of PCA graphs, but provided that one knows about this distortion, the gradient is still apparent: Both arches move clockwise from Individual 1 to 20 and from SNP 1 to 24. Also, those Individuals and SNPs that are located at similar positions along the gradient (in the obvious sense of having concentrations of 1s at the same position along the diagonal) are placed in similar directions from the origin—related Individuals and SNPs track each other in a clear and intuitive manner.

In the middle, the biplot shows the same gradient as at the left, but with the opposite option, SNP coding common=1. The arch for Individuals 1 to 20 is shown by black triangles, and the arch for SNPs 1 to 24 by green dots. Compared to Morrison’s original dataset, the arch for SNPs shown in green has been rotated by 180° relative to the Individuals shown in black. This rotation happens because reversal of the polarity of a SNP sends its point to the opposite location relative to the origin in a PCA graph—or more precisely, approximately to the opposite location because a low-dimensional graph approximates its high-dimensional data. Therefore, Individual 1 and SNP 1 are far apart with common=1, although they are near each other with rare=1 as they should be; and the same applies to Individual 20 and SNP 24. On the other hand, Individual 15 and SNP 8 nearly coincide with common=1 (near the horizontal dashed line, at the right), although they are far apart with rare=1 as they should be; and the same applies to Individual 6 and SNP 17 (at the left). Consequently, the biplot in the middle, unlike the original at the left, is counterintuitive and confusing because related Individuals and SNPs can be widely separated, and distant Individuals and SNPs can be quite close.

On the right, the biplot illustrates SNP coding mixed. In this dataset, the coding rare=1 and common=1 alternates, with columns using rare=1 shown in red, and those using common=1 shown in green. Incidentally, other mixed coding schemes give qualitatively the same results, such as selecting the coding at random for each SNP, or reversing the coding for every fourth SNP instead of every other SNP. The DC-PCA biplot on the right combines features of the left and middle biplots. Individuals 1 to 20 are shown with black triangles, odd-numbered SNPs 1 to 23 with red dots, and even-numbered SNPs 2 to 24 with green dots. The orientation of the green arch is rotated by 180° relative to the black and red arches. Taken together without distinguishing red from green, the dots for the SNPs roughly approximate a circle around the origin, rather than the typical arch, so awareness of the arch distortion would not be enough to guide proper interpretation. The biplot on the right inherits the problems from the middle biplot that related Individuals and SNPs can be widely separated, and distant Individuals and SNPs can be quite close. Furthermore, it has the additional problems that SNPs near each other along the gradient can be far apart in this biplot (such as SNPs 1 and 2), whereas SNPs far apart from each other along the gradient can be near each other (such as SNPs 4 and 15). Consequently, SNP coding mixed produces biplots that are quite confusing.

From Fig 4, the verdict on SNP codings for producing biplots is that SNP coding rare=1 is decidedly superior—invariably superior for mathematical reasons, rather than circumstantially superior for biological reasons. Furthermore, that verdict remains despite the arbitrary polarity of PCA components that is explained in Materials and Methods because arbitrary polarity applies to sign reversal of *both* row and column PCA scores, *not* just one or the other, so the orientation of the row arch and column arch relative to each other is invariant, not arbitrary.

However, the verdict on SNP codings for monoplots is different. Although the DC-PCA biplots in Fig 4 show markedly different results for the SNPs (red or green dots, or both), the results look identical for the Individuals (black triangles).

**Fig. 5.**
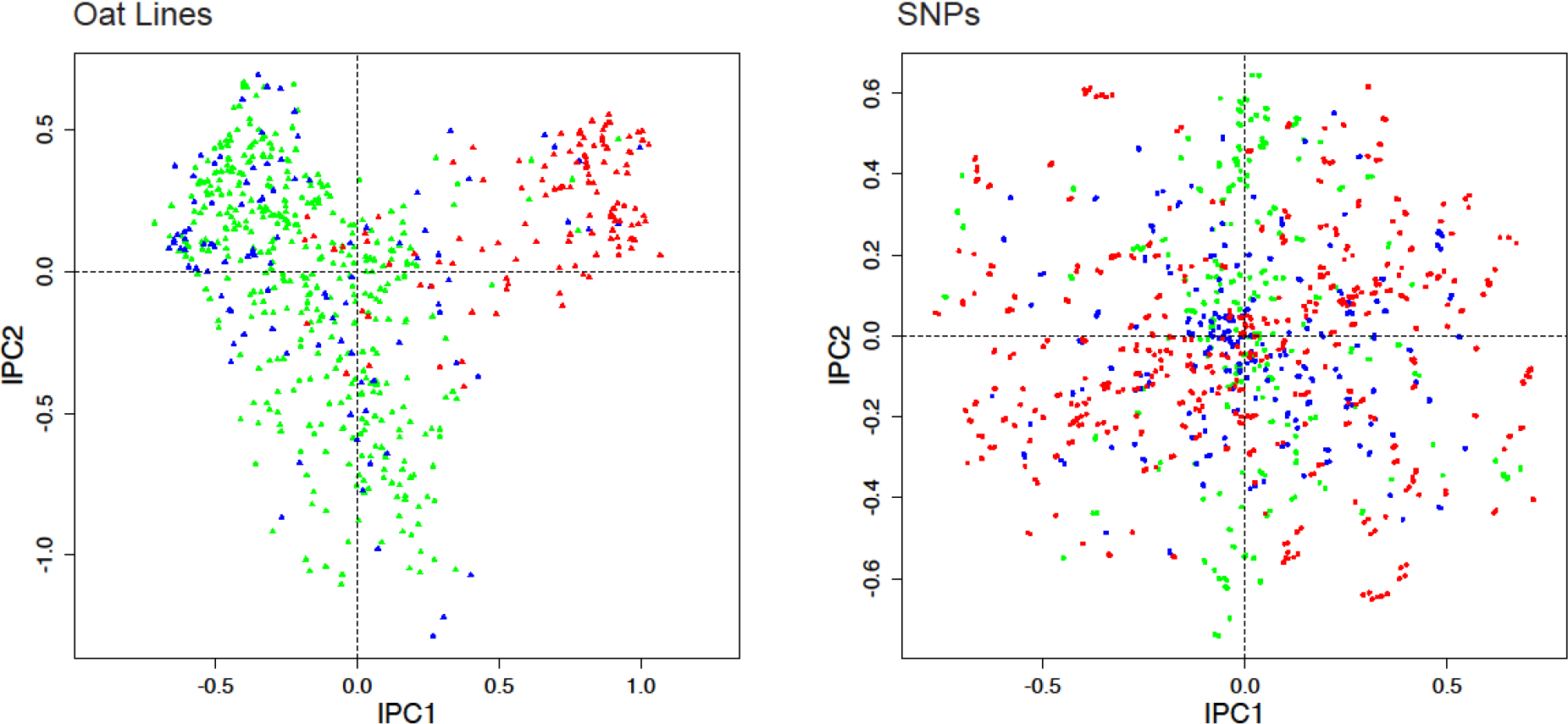
DC-PCA biplot for the oat data, using SNP coding mixed, namely the received data. The color scheme is the same as in Fig 1, namely spring oats show in green, world diversity oats in blue, and winter oats in red, with corresponding colors for the SNPs. The contrast between spring and winter oats is clear for Individuals in the left panel, but the corresponding pattern for SNPs in the right panel has been obliterated by thorough intermixing of the three colors of points.

Similarly, Fig 5 uses the oat dataset to reinforce principles learned about SNP codings with the toy dataset. The earlier Fig 1 used SNP coding rare=1. This figure substitutes SNP coding mixed, namely the original oat data as received from Kathy Esvelt Klos. As expected, the pattern for oat lines looks virtually the same in Figs 1 and 5. By contrast, the pattern for the SNPs in the right panel of Fig 5 is utterly obliterated, with no separation of green from red points. The explanation is that with the mixed coding of the received data, about half of the SNPs (772 out of 1341) have the reverse polarity common=1, which sends their points approximately to the opposite location in the DC-PCA biplot, and thereby thoroughly mixes the three colors of points for the SNPs. This outcome could be anticipated from the results for SNP coding mixed in Fig 4C. This example with real data might reinforce the suspicion that SNP coding is inconsequential for monoplots of Individuals only, even though SNP coding rare=1 is decidedly superior for biplots.

Nevertheless, although Figs 4 and 5 might suggest that SNP coding is inconsequential for the Individuals, actually it can matter. Fig 6 shows the biplot for the oat dataset using another SNP coding mixed, namely VCF coding using oat line 189 (a spring oat) as the reference genome. Of course, the pattern for the SNPs has been obliterated by coding mixed, but the pattern for oat lines has also changed noticeably from that in Fig 1. Furthermore, even for the toy dataset used in Fig 4, the term “look” the same was used to compare the patterns for Individuals in its three biplots, not “are” the same, because SNP coding mixed is very slightly different from rare=1 and common=1, although not enough different to be visually perceptible (as proven in the appendix). Likewise, the patterns for oat lines in Figs 1 and 5 look very similar, but careful inspection shows them to be slightly different.

**Fig 6.**
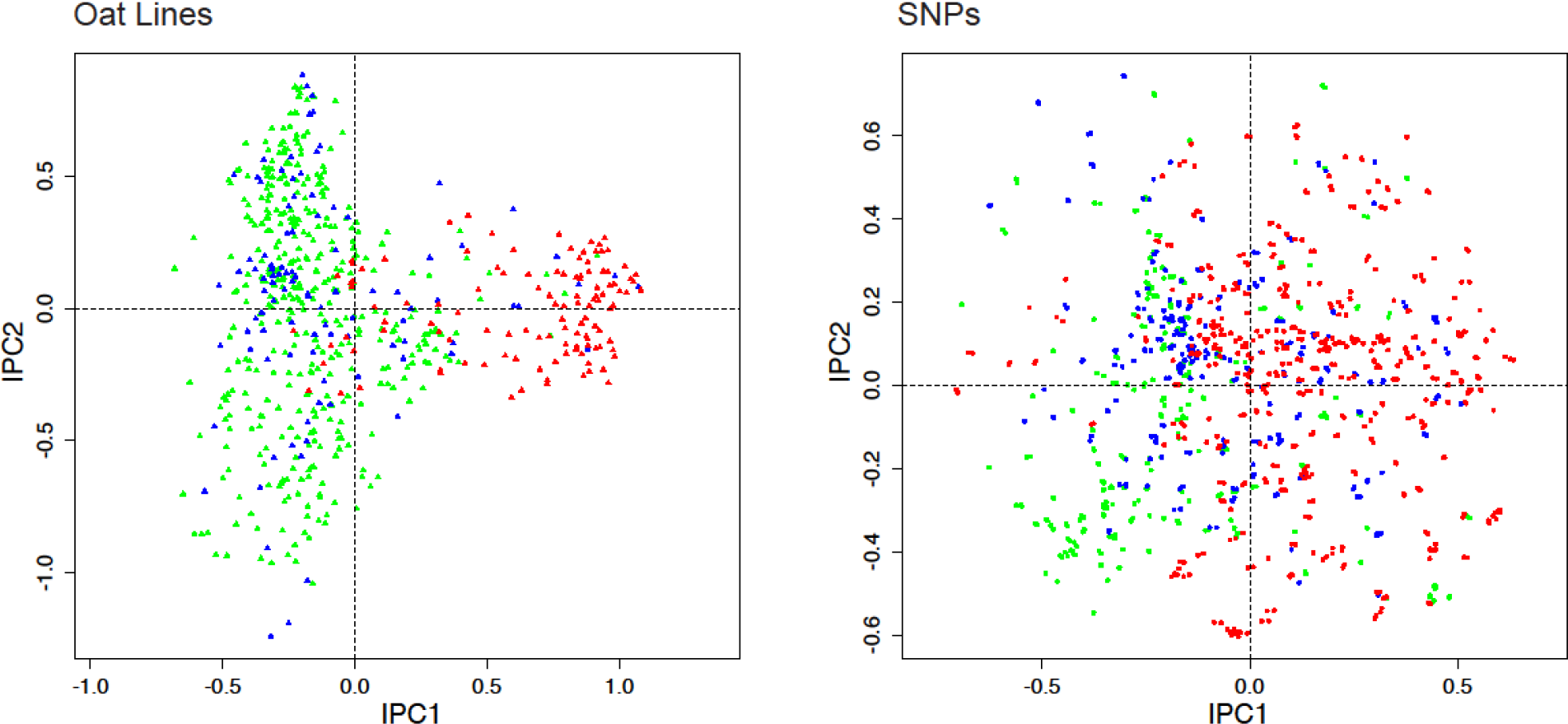
DC-PCA biplot for the oat data, using VCF coding with oat line 189 as the reference genome. The color scheme is the same as in Fig 1, namely spring oats show in green, world diversity oats in blue, and winter oats in red, with corresponding colors for the SNPs. Compared to Fig 1 with SNP coding rare=1, not only has the pattern for SNPs been totally obliterated, but also the pattern for oat lines has been changed noticeably.

The previous subsection mentioned *that* SNP coding rare=1 was used to construct the CA1 arranged matrix in Fig 2, and thereby to obtain the five colors used in Fig 3, whereas this subsection explains *why* SNP coding matters for CA. As ecologists have long known, although both PCA and CA have the arch distortion, PCA does but CA does not involute the arch, and therefore only CA can produce an arranged matrix that concentrates the larger values along the matrix diagonal (Figs 3.15 and 4.9 in [9]). But given the sort of data collected by ecologists to study plant and animal communities, naturally they have investigated only the sort of data represented in the present context by SNP coding rare=1. Actually, as illustrated for the same toy data used in Fig 4, indeed the CA1 arranged matrix does work for SNP coding rare=1 because its CA biplot is not involuted; but it does not work for SNP coding common=1 because its CA biplot is involuted, and it does not work for SNP coding mixed because its CA biplot is complicated (supplementary information S4 CA Results). Likewise, although SNP coding rare=1 enables CA1 order to display joint structure along the matrix diagonal for the oat data in Fig 2, both SNP coding common=1 and SNP coding mixed fail.

This subsection makes the important observation *that* SNP coding can affect PCA results for Individuals, whereas the next subsection and appendix provide the explanation for *why* SNP coding can matter, even for researchers who are interested only in results for Individuals. Furthermore, although Fig 6 happens to illustrate a modest change in the pattern for Individuals, depending on circumstances, choices among SNP codings can also cause drastic changes.

In review, the three choices of SNP coding, namely rare=1 or common=1 or mixed, are often inconsequential for DC-PCA monoplots of only the Individuals, but not always. However, choices of SNP coding are hugely consequential for DC-PCA biplots of both Individuals and SNPs, for which rare=1 is far superior for displaying structure or patterns in the SNPs in a manner that is readily interpreted.

### Choices between six PCA variants

The biological information displayed in the recommended DC-PCA biplot has exceptional clarity and straightforward interpretation because DC-PCA uniquely avoids confounding of main and interaction effects, and thereby it avoids multiple and complicated interpretive principles that necessitate a special augmented ANOVA table in order to discern for each individual dataset what sort of information is presented in its biplot. Again, the six PCA variants investigated here are SNP-Centered, Individual-Centered, Double-Centered, SNP-Standardized, Individual-Standardized, and Grand-Mean-Centered PCA. The two choices of the first subsection, monoplots or biplots, exhaust the possibilities; and the three choices of the second subsection, SNP codings rare=1 or common=1 or mixed, also exhaust the possibilities. However, the six choices of this third subsection do not exhaust the possibilities, so this subsection offers a useful but not comprehensive account of data transformations and hence PCA variants.

The statistical meanings of the three sources of variation—SNP main effects, Individuals main effects, and S×I interaction effects—are fundamentally different. Consider a data matrix with *p* SNPs and *n* Individuals. The SNP main effects concern the means across Individuals for each SNP, so they constitute a vector of length *p*. Likewise, the Individual main effects concern the means across SNPs for each Individual, so they constitute a vector of length *n*. By contrast the S×I interactions equal the data minus both main effects, so they constitute a matrix of dimensions *p* and *n*. Main effects are relatively simple and can easily be tabulated or graphed by a variety of familiar methods, whereas interaction effects are complex and require multivariate statistical analyses such as PCA. These three sources are radically different statistically in the strong sense that they are orthogonal and uncorrelated, so knowing any one of them provides no information whatsoever on the other two.

The biological meanings of these three sources are as follows in the present context of a SNPs-by-Individuals data matrix. For a given SNP, its mean across Individuals is simply the frequency of the allele coded 1 (presuming that the alternative allele is coded 0). Likewise, for a given Individual, its mean across SNPs is simply the frequency of the allele coded 1. Ordinarily, it makes sense to avoid burdening or distracting a PCA graph with such simple information, so it is best to remove main effects. Also, these means often lack any straightforward or interesting biological meaning. Indeed, it may be especially difficult to attach any biological meaning to the Individual means, not least because of alternative choices for SNP coding. When SNP or Individual means are biologically meaningless, they merely add noise to a PCA graph.

That said, occasionally there can be special circumstances for which main effects are also of interest, in addition to the S×I effects. Which of these three sources of variation—SNP, Individual, and S×I effects—are of interest depends on the data and the research objectives. It is the prerogative of researchers to decide which of these sources interest them. Our default recommendation is DC-PCA, which focuses on S×I exclusively; but we oblige researchers with a different focus by exploring additional variants of PCA in this subsection.

A geometrical interpretation of PCA can convey an intuitive grasp of the unique conceptual appropriateness of DC-PCA for elucidating population structure. Consider a data matrix with *R* rows and *C* columns. These data can be conceptualized as a high-dimensional cloud of *R* points in *C*-dimensional space, or conversely as *C* points in *R*-dimensional space (Table 4.1 and Fig 4.1 in [9]). DC-PCA amounts to translation of the coordinate system to the centroid of the cloud of points, followed by rigid rotation of the coordinate axes such that perpendicular projections of the points onto PC1 maximizes the variance captured by this axis, then PC2 captures the most remaining variance, and so on for higher PCs (Fig 4.6 in [9]). The origin for the PCA rotation is located in high-dimensional space, and in all of our biplots it is projected onto the PC1-PC2 plane at the intersection of the dashed horizontal and vertical lines located at (0, 0). Insofar as the centroid is the eminently sensible place to put the origin, DC-PCA makes sense because it uniquely places the origin at the centroid of the cloud of points in both panels of a biplot (as in Figs 1 and 3, or for both triangles and dots in Fig 4). By contrast, some published PCA monoplots of Individuals, which show population structure with variants other than DC-PCA, have a unipolar PC1 (all positive or all negative scores, rather than both which includes 0) so extreme that rotation of the coordinate system to PCs occurs at an origin located outside the graph. In such cases, one would expect special reasons to be given to justify such a peculiar location of the origin, and one can only wonder what the population structure would have looked like were the PCA rotation around the centroid, which ordinarily makes much more sense. As explained later in this subsection, dislocating the origin of the PCA away from the centroid of the data equates to confounding main and interaction effects, which complicates interpretation of population structure in ways that the genomics community does not yet realize whenever PCA graphs are made with variants other than the recommended DC-PCA.

Table 1 shows the ANOVA table for DC-PCA of the oat data using three SNP codings: rare=1, VCF with oat line 189 as the reference genome (which is of particular interest because it has larger Individual main effects than any of the other 634 possibilities), and the SNP coding for the data as received (and shifted to 0 and 1). ANOVA partitions the total degrees of freedom (df) and sum of squares (SS) into three sources, and then PCA partitions the S×I interactions into the first seven IPCs followed by the residual. The sources are indented to indicate subtotals.

**Table 1.** ANOVA table for DC-PCA of SNP data on oats using three SNP codings: rare=1 (and common=1 is identical), VCF with oat line 189 as the reference genome, and the data as received. Both VCF189 and the received data are instances of SNP coding mixed.

| Source | df | SS rare=1 | SS VCF189 | SS received |
| --- | --- | --- | --- | --- |
| Total | 851534 | 157442.756 | 175776.962 | 210084.641 |
| SNPs | 1340 | 16872.731 | 35206.937 | 69514.616 |
| Individuals | 634 | 2077.808 | 6286.978 | 482.151 |
| S×I | 849560 | 138492.217 | 134283.047 | 140087.874 |
| IPC1 | 1973 | 15566.723 | 12202.771 | 16158.469 |
| IPC2 | 1971 | 9512.804 | 9172.430 | 9687.670 |
| IPC3 | 1969 | 5836.915 | 6163.309 | 6246.232 |
| IPC4 | 1967 | 4831.287 | 4140.115 | 4803.660 |
| IPC5 | 1965 | 3507.237 | 3478.698 | 3488.106 |
| IPC6 | 1963 | 2997.351 | 3100.062 | 3181.724 |
| IPC7 | 1961 | 2887.575 | 2839.061 | 2881.938 |
| Residual | 835791 | 93352.327 | 93186.601 | 93640.076 |

For SNP coding rare=1, the total SS is composed of 88.0% for S×I interaction effects, 10.7% for SNP main effects, and 1.3% for Individual main effects. SNP coding common=1 is not shown in Table 1, but it necessarily has exactly the same ANOVA table as SNP coding rare=1, not only for DC-PCA shown here, but also for all six variants of PCA considered in this article. However, VCF for oat 189 has different percentages, namely 76.4%, 20.0%, and 3.6%, and the received data has 66.7%, 33.1%, and 0.2%. Hence, choices of SNP coding affect the relative magnitudes of these three sources, as well as the relative magnitudes of the IPCs.

The application of PCA to a combination of two sources of variation, unlike the single source of S×I interaction effects for DC-PCA in Table 1, requires a new approach in order to understand what kind of information is in each PC, namely an augmented ANOVA table that is introduced here for the first time. The SS of each PC is partitioned into the portions due to main and interaction effects. The required calculations are simple: For each PC, multiply its SNP scores and Individual scores, which are a row vector and a column vector, to obtain the matrix of expected values, and then subject that matrix to ANOVA. Because augmented ANOVA tables are not yet included in available software, we wrote our own R code (supporting information S5 Software). Table 2 shows an augmented ANOVA table for SNP-Centered PCA of the same oat data, using SNP coding rare=1. This variant of PCA removes only SNP main effects, and then applies PCA to the Individual main effects *and* S×I interaction effects combined, denoted by I&S×I, which has a SS of 2077.808 + 138492.217 = 140570.025. Researchers who are familiar with PCA and are accustomed to the automatic monotonic decrease in the SSs for successive PCs should note that the SSs for the Individuals and S×I portions are not necessarily monotonic.

**Table 2.** Augmented ANOVA table for SNP-Centered PCA of SNP data on oats, using SNP coding rare=1. PCA is applied to Individual main effects and S×I interaction effects combined (I&S×I), and the portion of each is shown in the last two columns.

| Source | df | SS | Individuals | S×I |
| --- | --- | --- | --- | --- |
| Total | 851534 | 157442.756 |  |  |
| SNPs | 1340 | 16872.731 |  |  |
| I&S×I | 850194 | 140570.025 | 2077.808 | 138492.217 |
| PC1 | 1974 | 16325.612 | 857.676 | 15467.936 |
| PC2 | 1972 | 9751.158 | 257.217 | 9493.941 |
| PC3 | 1970 | 6250.504 | 423.134 | 5827.370 |
| PC4 | 1968 | 4860.910 | 23.386 | 4837.524 |
| PC5 | 1966 | 3510.312 | 4.521 | 3505.791 |
| PC6 | 1964 | 3181.953 | 187.615 | 2994.338 |
| PC7 | 1962 | 2891.656 | 1.654 | 2890.002 |
| Residual | 836418 | 93797.920 | 322.606 | 93475.314 |

All seven PCs and the residual of SNP-Centered PCA contain a mixture of Individual and S×I effects. Such mixtures always occur for any dataset whenever PCA is applied to a combination of main and interaction effects [20]. For this particular dataset, the first seven PCs and the residual are all dominated by S×I interaction effects because the Individual main effects happen to be small. That outcome could be expected from Table 1 since IPC1 through IPC7 are all larger than the Individual main effects. Comparing the ANOVA for SNP coding rare=1 in Table 1 with the augmented ANOVA in Table 2, PC1 and PC2 from SNP-Centered PCA are larger than IPC1 and IPC2 from DC-PCA because main effects for Individuals are confounded with S×I interaction effects, but more importantly, IPC1 and IPC2 for DC-PCA capture *more* S×I interactions than PC1 and PC2 from SNP-Centered PCA because DC-PCA is not distracted by any main effects. This comparison can be understood in geometrical terms by saying that obviously without centering PCA is wasting effort to capture the non-central centroid of the data, and that distraction compromizes PCA’s visualization of the data.

SNP-Centered PCA has four possible outcomes. The oat example in Table 2 illustrates one possibility, that S×I information dominates both PC1 and PC2. Indeed, the Individual main effects account for only 4.3% of the SS captured in a PC1-PC2 graph. Another possible outcome, caused by main effects having a larger SS than does PC1, is that PC1 contains mostly main-effect information whereas PC2 contains mostly S×I information. Yet another possible outcome, caused by main effects having a larger SS than PC2 but a smaller SS than PC1, is the reverse, that PC1 contains mostly S×I information and PC2 contains mostly main-effect information. Finally, it is also possible for a PC to contain substantial portions of both main and interaction effects. For example, SNP-Centered PCA using VCF coding with oat line 189 as the reference genome has a PC1 comprised of 31.1% Individual main effects and 68.9% S×I interaction effects (Table S6.1 in supplementary information S6 Four Tables). It is crucial for researchers to know which of these four cases obtains for a given dataset when they interpret a PC1-PC2 graph that uses SNP-Centered PCA.

Individual-Centered PCA also has four possible outcomes. Table 3 shows the augmented ANOVA table for Individual-Centered PCA of the same oat data, using SNP coding rare=1. This variant of PCA removes Individual main effects and then applies PCA to the SNP main effects and S×I interaction effects combined, denoted by S&S×I, which has a SS of 16872.732 + 138492.217 = 155364.949. The table shows that PC1 is dominated by SNP main effects (96.2%), whereas PC2 is dominated by S×I interaction effects (99.9%). That outcome could be expected from Table 1 since the SNP main effects are larger than IPC1. As already explained for SNP-Centered PCA in Table 2, this example illustrates only one of the four possible outcomes.

**Table 3.** Augmented ANOVA table for Individual-Centered PCA of SNP data on oats, using SNP coding rare=1. PCA is applied to SNP main effects and S×I interaction effects combined (S&S×I), and the portion of each is shown in the last two columns.

| Source | df | SS | SNPs | S×I |
| --- | --- | --- | --- | --- |
| Total | 851534 | 157442.756 |  |  |
| Individuals | 634 | 2077.808 |  |  |
| S&S×I | 850900 | 155364.948 | 16872.731 | 138492.217 |
| PC1 | 1974 | 17402.420 | 16734.514 | 667.906 |
| PC2 | 1972 | 15564.018 | 21.919 | 15542.099 |
| PC3 | 1970 | 9476.489 | 40.283 | 9436.206 |
| PC4 | 1968 | 5781.809 | 23.612 | 5758.197 |
| PC5 | 1966 | 4758.747 | 25.746 | 4733.001 |
| PC6 | 1964 | 3482.969 | 5.452 | 3477.517 |
| PC7 | 1962 | 2970.194 | 4.894 | 2965.270 |
| Residual | 837124 | 95928.303 | 16.312 | 95912.021 |

Like their centered counterparts, PCs from SNP-Standardized and Individual-Standardized PCA contain a mixture of main and interaction effects, so these PCA variants also have four possible outcomes. Furthermore, SNP-Standardized PCA, unlike SNP-Centered PCA, cannot be used with VCF coding because the reference genome has a standard deviation of zero.

Grand-Mean-Centered PCA produces a mixture of SNP main effects, Individual main effects, and S×I interaction effects in each PC. Therefore, the situation for Grand-Mean-Centered PCA is quite complex and undesirable: It has seven possibilities, not counting additional possibilities involving a PC with a substantial mixture of main and interaction effects. The supporting information includes the augmented ANOVA tables for these additional variants, using the oat data with SNP coding rare=1 (Tables S6.2 to S6.4 in S6 Four Tables). Between the main text and the supporting information, ANOVA tables are shown for all six PCA variants.

When main effects are of interest, we recommend the Additive Main effects and Multiplicative Interaction (AMMI) model, which combines ANOVA for the main effects with PCA for the multiplicative effects [20]. AMMI and DC-PCA are similar and have an identical ANOVA table. The salient difference is that whereas DC-PCA discards the main effects, AMMI retains them. An AMMI1 biplot shows both of the main effects in its abscissa, and IPC1 in its ordinate; it can show only one component (and hence the suffix 1 in AMMI1) because the main effects use one of its two axes. Its abscissa captures 100% of both main effects. Also, its ordinate captures as much of the S×I interaction effects as possible because IPC1 is the unique least-squares solution that maximizes the variation along this axis and minimizes the residual variation off this axis. We have not yet encountered AMMI in genomics, but it is commonplace in the literature on agricultural yield trials [20]. Incidentally, in statistical analyses of agricultural yield trials, the so-called AMMI2 biplot shows IPC1 and IPC2, which is exactly what a DC-PCA biplot (ordinarily) shows, so “AMMI2” and “DC-PCA” are two names used in different literatures for the same analysis.

Fig 7 shows the AMMI1 biplot for the oat data, using SNP coding rare=1 and the same color scheme as Fig 1. The abscissa shows the mean frequency of the rare allele. The oat lines have means that range from 0.15958 to 0.44893, and the SNPs range from 0.01732 to 0.49921. The vertical line is located at the grand mean of 0.24484. The abscissa captures 100% of the SNP main effects, 100% of the oat line main effects, and 0% of the S×I interaction effects. The ordinate of Fig 7 shows IPC1, which captures 11.2% of S×I, and this ordinate is identical to the abscissa in Fig 1; as before IPC1 separates green (spring oats) from red (winter oats) for both oat lines and SNPs. In the right panel, the SNPs have a broad range of IPC1 scores at the right but not at the left because SNPs at the right have large numbers of both rare and common alleles and hence have large variances, whereas SNPs at the left have mostly the common allele and hence have small variances.

**Fig 7.**
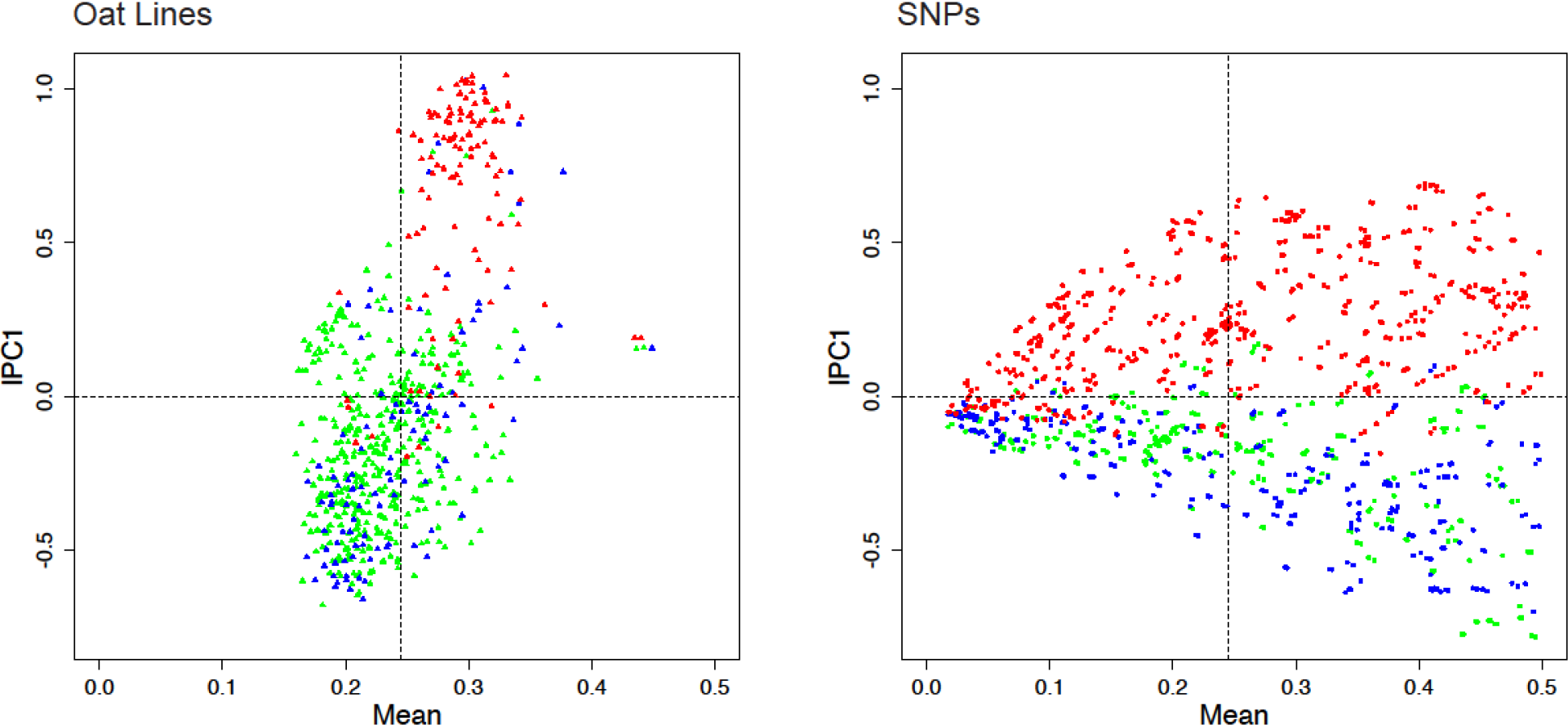
The AMMI1 biplot for the oat data, using SNP coding rare=1. To reduce clutter the biplot uses two panels, with oat lines on the left and SNPs on the right. The color scheme is the same as in Fig 1, namely spring oats show in green, world diversity oats in blue, and winter oats in red, with corresponding colors for the SNPs.

The interpretive principles for an AMMI1 biplot are that displacements along the abscissa reflect differences in main effects, whereas displacements along the ordinate reflect differences in S×I interaction effects. An Individual and SNP with IPC1 scores of the same sign have a positive S×I interaction, whereas those with opposite signs have a negative S×I interaction, and an Individual (or SNP) with a score near zero has small interactions with all SNPs (or Individuals)—at least for those interactions that are captured in the AMMI1 biplot. Like DC-PCA, because AMMI1 has a single kind of information in its abscissa and in its ordinate, an AMMI1 biplot has a single set of interpretive principles that applies to all datasets.

A researcher may make a choice between not having, or else having, interest in main effects—which equates to the difference between DC-PCA and the other five variants of PCA considered here. However, the existence of AMMI provides a *further choice* between *confounded* main and interaction effects from PCA variants other than DC-PCA, including SNP-Centered PCA, or else *non-confounded* main and interaction effects from AMMI. Given this further choice between confounded or non-confounded effects, we are not aware of any plausible arguments in favor of confounded effects. Accordingly, we recommend DC-PCA or AMMI1 to accommodate researchers’ diverse interests, but disfavor all of the other five variants of PCA because of confounding and the consequent multiple outcomes for how their biplots should be interpreted.

Five additional PCA biplots for the oat dataset are shown in the supporting information, using SNP coding rare=1 and the same color scheme as the main text (S7 Five Biplots). SNP-Centered and SNP-Standardized PCA approximate DC-PCA in Fig 1 because the Individual main effects are small, whereas Individual-Centered, Individual-Standardized, and Grand-Mean-Centered PCA approximate AMMI1 in Fig 7 because IPC1 captures mostly the large SNP main effects. The biplots for Individual-Centered and Grand-Mean-Centered PCA have a unipolar PC1 in the panel for oat lines. However, although these other PCA variants can approximate DC-PCA or AMMI1, they can *only* approximate because every component has a mixture of main and interaction effects. Between Fig 1 the main text and the supporting information, biplots are shown for all six PCA variants, with all six biplots using SNP coding rare=1 and the same color scheme as in Fig 1.

The findings from ANOVA tables in this subsection reinforce the value of biplots that was stressed in the first subsection. The joint structure of SNPs-and-Individuals just is the S×I interactions, which only biplots can display, and S×I is commonly large. From Table 1, the oat data contains 88.0% interaction information (138492.217 / 157442.756), using SNP coding rare=1. Four additional examples can be cited for other crop species, using whatever SNP coding the original authors selected: rice (*Oryza sativa* L.) [21] has 91.5% interaction information, soybean (*Glycine max* (L.) Merr.) [22] has 88.7%, maize (*Zea mays* L.) [23] has 65.2%, and potato (*Solanum tuberosum* L.) [24] has 30.1%. Furthermore, the percentage of interaction information captured in a PC1-PC2 graph is often higher than that in a dataset as a whole, as quantified by an augmented ANOVA table. For instance, the percentages of interaction in a PC1-PC2 biplot for SNP-Centered PCA for these five species in the same order are 95.72%, 98.34%, 94.49%, 73.73%, and 99.93%.

In review, both SNP coding and PCA variant can affect which kind of information— Individual or SNP or S×I effects—dominates in each PC for all variants of PCA other than DC-PCA. An augmented ANOVA table quantifies the outcome for any dataset, and thereby facilitates proper interpretation of PCA results and graphs. Our default recommendation is DC-PCA because it uniquely applies PCA to a single source of variation, namely the S×I interactions that are often of primary biological interest, so its IPCs always contain this one kind of information and hence there is no need for its ANOVA table to be augmented. When main effects are of interest, we recommend an AMMI1 biplot because it uniquely displays main and interaction effects without confounding them.

## Discussion

Because PCA monoplots of only Individuals provide some insight into population structure, they are deservedly popular in the literature. This article can explain that success. Because only Individuals are graphed, unknowingly but luckily the choice of SNP coding is often inconsequential. And because the SS for Individual main effects is often small relative to the SS for IPC1, unknowingly but luckily some other PCA variants, namely SNP-Centered and SNP-Standardized PCA, may approximate the recommended DC-PCA. However, this success has serious limitations. The obvious problem is that relying on luck to obtain a usable monoplot of Individuals is not as reliable as deliberately choosing the recommended SNP coding rare=1 and PCA variant DC-PCA. The main problem is that a monoplot cannot show interaction structure, which is often the dominant source of variation in a dataset and is usually the variation of principal interest. Production of a useful biplot is an unlikely prospect apart from understanding the consequences of SNP codings and PCA variants.

About 80% of PCA graphs in the SNP literature show an evident PCA arch, but regrettably nearly 0% of these articles mention this distortion and explicitly take it into account when interpreting PCA graphs. We agree with Morrison’s blogposts that it would be good for this situation to change. Clearly, an assumption that the readership of these articles is already well aware of the PCA arch distortion is unwarranted. Consequently, there has been a missed opportunity to note and properly interpret a PCA arch (which appears as a two-dimensional structure in a PC1-PC2 graph) as a single trend following along the arch—a single trend that can often be related to a single underlying causal factor or gradient. There has also been a missed opportunity to perceive a PCA arch as an invitation to construct graphics such as Figs 2 and 3 in order to see whether they happen to reveal, for a given SNP dataset and research purpose, additional data structure of biological interest.

Elucidating population structure helps to accomplish many other research purposes. For example, population structure is important for genome-wide association studies (GWAS) because the first few PCs can be used as covariates in a regression to address the problem of spurious associations produced by population structure [13]. In turn, population structure and GWAS are important for plant breeding and human medicine [25].

This article addresses just one statistical analysis, PCA, applied to just one kind of genomics data, SNPs, for just one research interest, elucidating population structure. But this is just the tip of the iceberg: The genetics and genomics community uses many statistical analyses for many kinds of data and many research interests. Indeed, PCA graphs of SNP data commonly occur in multi-panel figures that also show neighbor-joining trees and clustering by Bayesian or other methods, as well as geographical or other relevant information [26–28], and a CA1 arranged matrix is yet another graphic that such figures could include. Given the opportunities to improve upon contemporary practices for PCA analysis of SNP data, it would not be too surprising if a wider exploration of statistical analyses of genomics data discovered additional opportunities to increase research productivity.

Finally, perhaps the most significant and promising result from this investigation is the opportunity for researchers to ask new questions that have not been asked before: What population structure has been present in SNP datasets all along, which cannot be displayed by contemporary practices of PCA analysis, but can be elucidated by the recommended choices of PCA graph, SNP coding, and PCA variant? And how vital is that additional biological insight for achieving research objectives? Answers to these questions can emerge only from the accumulated experiences of many researchers working with many SNP datasets and diverse research objectives. What can be concluded already, however, is that necessarily and unavoidably, *every* PCA graph of SNP data requires and implements *some* choices of PCA graph, SNP coding, and PCA variant—whether or not these fateful choices are reported and justified. And having already collected SNP data and produced a PCA graph with whatever choices, very little additional effort would be required to repeat the PCA analysis with the three choices recommended here, and then to discern what additional population structure is displayed and discovered.

## Conclusions

Three principal recommendations emerge from this investigation into PCA analysis of SNP data. (1) Use biplots, not monoplots, since only they can display S×I interaction information, that is, the joint structure of SNPs-and-Individuals. (2) Use the SNP coding 1 for the rare allele and 0 for the common allele. (3) Use the PCA variant DC-PCA if only S×I interactions are of interest, as is often the case; otherwise, use AMMI1 if main effects are also of interest. Additionally, report which SNP coding and PCA variant were selected, and ideally also provide reasons for those particular choices, so that readers can interpret PCA results properly and reproduce PCA analyses reliably. Finally, if the recommended DC-PCA (or AMMI1) is not used, then provide an augmented ANOVA table in order to quantify the amount of main and interaction effects in each PC. These conclusions should be understood as useful default recommendations because some dataset properties or research purposes may constitute exceptions, such as a focused research interest in only the Individuals that calls for publication of a monoplot—although it may still be worthwhile to produce the biplot and inspect it carefully before making a final decision about which graph to publish. An understanding of the consequences of choices between two PCA graphs, three SNP codings, and six PCA variants is an asset for elucidating population structure.

## Materials and methods

### Construction and interpretation of PCA graphs

A convenient form of the equation for DC-PCA is:
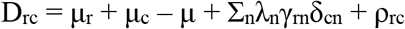
where D_rc_ is the datum for row *r* and column *c*, μ_r_ is the mean for row *r*, μ_c_ is the mean for column *c*, μ is the grand mean, λ_n_ is the singular value for component *n* and λ_n_^2^ is its eigenvalue, γ_rn_ is the eigenvector value for row *r* and component *n*, δ_cn_ is the eigenvector value for column *c* and component *n*, with both eigenvectors scaled as unit vectors, and ρ_rc_ is the residual for row *r* and column *c* when the number of components used is fewer than the full model (namely, one less than the minimum of the number of rows and number of columns). The singular values and eigenvectors are obtained by SVD, as explained in the appendix.

The convention adopted here is to multiply eigenvector values by the square root of the singular value to obtain λ_n_^0.5^γ_rn_ and λ_n_^0.5^δ_cn_ so that their products estimate row-by-column interactions directly without need for another multiplication by λ, and to call these values “PCA scores.” The PCA literature does not have consistent terminology since sometimes the results for rows (or individuals in the present context) are called “PCA scores” whereas results for columns (or SNPs) are called “PCA loadings.” Because PCA is symmetric regarding matrix rows and columns, as the above equation makes evident, we prefer to use the single and consistent terminology of PCA scores. However, other contexts may call for a different approach, such as a PCA monoplot of Individuals, for which multiplication by the singular value is preferable because this optimizes the two-dimensional approximation of distances between Individuals. For a comprehensive explanation of ways to handle the singular value, see Malik and Piepho [29].

The axes in PCA graphs are often scaled to obtain a convenient shape, but actually the axes should have the same scale for many reasons [29]. Unfortunately, our literature survey found a correct ratio of 1 in only 10% of the articles, a slightly faulty ratio of the larger scale over the shorter scale within 1.1 in 12%, and a substantially faulty ratio above 2 in 16%, with the worst cases being ratios of 31 and 44. Also, 7% of the articles failed to show the scale on one or both PCA axes. However, the two axes of an AMMI1 biplot contain different kinds of information (main or interaction effects), so they do *not* need to use the same scale.

The solution to SVD is unique, up to simultaneous sign change of both eigenvectors for a given component. This is evident from the above PCA equation because the terms in the equation λ_n_γ_rn_δ_cn_ are products of the two scores λ_n_^0.5^γ_rn_ and λ_n_^0.5^δ_cn_, so reversing the signs of *both* scores leaves their products unchanged. Because of arbitrary polarity, different software may produce PCA analyses for a given dataset with opposite polarity for some or all components. This has two implications for producing and interpreting PCA graphs, including biplots. First and most important, polarity reversal leaves all distances between points unchanged, so mathematically and geometrically, it is absolutely inconsequential. Indeed, reversing the polarity of an ordinate amounts to flipping a graph over from left to right, or reversing the polarity of an abscissa amounts to flipping a graph over from top to bottom, and reversing both axes amounts to rotating a graph by 180°—nothing changes. Second, especially when several PCA graphs of the same or related data are being compared with each other, adopting a consistent orientation constitutes good pedagogy, making it easier for readers to perceive the salient differences between graphs without distraction from arbitrary orientations.

The PCA arch or horseshoe distortion is important because it generates both problems and opportunities as explained in the Results section, and also because it occurs in the PC1-PC2 plane, which is what researchers show most often in PCA graphs. But this distortion, which amounts to a quadratic representation of PC1 in PC2, is only the first of a polynomial sequence. Using simulated genomics data, graphs of PC1 through PC4 on the abscissa and a one-dimensional habitat or environmental gradient on the ordinate show a sequence of linear, quadratic, cubic, and quartic polynomials [13]. The same was shown three decades earlier in the context of ecology using a related statistical analysis, CA (Fig 4 in [30]). These additional polynomials must be recognized when interpreting graphs other than the basic PC1-PC2 graphs, such as the three dimensional PC1-PC2-PC3 graphs and PC1-PC3 or PC2-PC3 graphs that appear in the genomics literature occasionally. Special graphics can show population structure for many components, such as PC1-PC12 (Fig 3 in [31]). The PCA arch in the first two components, and the polynomial sequence in higher components, is a feature of not only PCA and CA, but also NMS [32].

The following principles for interpreting biplots pertain specifically to DC-PCA. For two points of the same kind (two Individuals or else two SNPs), nearby points have similar interactions, whereas distant points have dissimilar interactions. For two points of different kinds (an Individual and a SNP), Individuals in a given direction from the origin have positive interactions for SNPs in that same direction, Individuals in a given direction have negative interactions for SNPs in the opposite direction, and Individuals in a given direction have small interactions for SNPs at right angles. Accordingly, interactions are large when an Individual and SNP are both far from the origin (but not at right angles), whereas an Individual near the origin has small interactions with all SNPs, and likewise a SNP near the origin has small interactions with all Individuals. More exactly, these interpretive principles address the interaction structure captured in the IPC1-IPC2 plane of a biplot.

The percentage of variation captured by each PC is often included in the axis labels of PCA graphs. In general this information is worth including, but there are two qualifications. First, these percentages need to be interpreted relative to the size of the data matrix because large datasets can capture a small percentage and yet still be effective. For example, for a large dataset with over 107,000 SNPs and over 6,000 persons, the first two components capture only 0.3693% and 0.117% of the variation, and yet the PCA graph shows clear structure (Fig 1A in [33]). Contrariwise, a PCA graph could capture a large percentage of the total variation, even 50% or more, but that would not guarantee that it will show evident structure in the data. Second, the interpretation of these percentages depends on the choice of a PCA variant, as augmented ANOVA tables make clear. Readers cannot meaningfully interpret the percentages of variation captured by PCA axes when authors fail to communicate which variant of PCA was used.

The objection may be raised that PCA biplots would be impractical for datasets with many thousands of SNPs, making graphs unworkably cluttered. In fact, high-density PCA graphs appear in the literature routinely, such as Fig 4 in [34] showing results for 54734 humans. Producing biplots in two adjacent panels helps to reduce clutter by separating Individuals from SNPs. Fortunately, the literature offers several strategies for simplifying PCA graphs. One possibility is to reduce the number of SNPs prior to PCA, using tools such as PLINK [35] and bigstatsr or bigsnpr [36]. Another is to select SNPs of particular interest. For example, only 23 SNPs out of over 1,000,000 produce PCA graphs with clear clusters for several major US populations [37, 38]. Obviously, it is impossible to label thousands of points without causing severe overprinting, but when only a moderate number of Individuals or SNPs are of special interest, they can be labeled.

Enormous SNP datasets are becoming increasingly common, and fast PCA algorithms can readily handle large-scale genome-wide data. The remarkably efficient software FastPCA computes the top several PCs with time and memory costs that are linear in the number of matrix entries [34]. The software flashpca is also very fast [39]. The power method is the simplest algorithm for PCA and is efficient when only the first few PCs are needed [40]. Incidentally, the power method is simple and fast for obtaining just the first component of CA (Appendix 2 in [15]), with time and memory costs that are linear in the number of matrix entries.

### Literature survey

The 125 articles applying PCA analysis to SNP data were taken from the literature more or less at random, with some emphasis on agricultural crop species and on researchers at Cornell University. They span many species and many journals. This survey is included in the supporting information (S1 Literature Survey).

### Oat datasets

The oat dataset supplied by Kathy Esvelt Klos is included in two formats: SNP coding mixed is the data as received, except that the original coding of 1 and 2 was shifted to 0 and 1; and SNP coding rare=1, which required polarity reversal for 772 of the 1341 SNPs (S2 Oat Mixed and S3 Oat Rare1).

### PCA and CA analyses

Our R code for comparing six PCA variants and correspondence analysis (CA) is included in the supporting information (S5 Software). From the R library, our code uses ca for CA and ggplot2 for graphs.

## Supporting information

S1 Literature Survey

S2 Oat Mixed

S3 Oat Rare1

S4 CA Results

S5 Software

S6 Four Tables

S7 Five Biplots

## Appendix: Consequences of SNP coding for six variants of PCA

This appendix concerns which variants of PCA are, or else are not, immune to changes in SNP coding as regards PCA monoplots of Individuals, where “Individuals” is a generic term for samples such as persons or cultivars. The main text already showed in Table 1 that SNP coding affects the sum of squares (SS) for SNP main effects. Therefore, Individual-Centered PCA is not immune because different proportions of main and interaction effects can change which PC is dominated by the SNP main effects, thereby dramatically altering a PCA monoplot of Individuals. This same verdict of not being immune also applies to Individual-Standardized PCA for the same sort of reason. Likewise, Grand-Mean-Centered PCA is not immune because it also retains SNP main effects (and Individual main effects), and again SNP coding affects the SS for SNP main effects. The remainder of this appendix addresses the remaining three variants in the order SNP-Centered, SNP-Standardized, and Double-Centered PCA.

First, consider SNP-Centered PCA. Let *Y* be the *p* × *n* SNP data matrix with SNPs in *p* rows and Individuals in *n* columns. Without loss of generality, assume that *p*≥ *n*. The matrix *Y* may be SNP-Centered as follows: 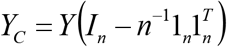, where *I*_*n*_ is the *n*-dimensional identity matrix and 1_*n*_ is an *n*-vector of ones. Let *Y*_*c*_=*USV*^*T*^ be a singular value decomposition of *Y*_*c*_, where *U* is a *p* × *n* orthonormal matrix of left singular vectors holding the row scores, *V* is an *n* × *n* orthogonal matrix right singular vector holding the column scores, and *S* is a diagonal matrix of order *n* holding the ordered singular values. From the orthonormality of *U* we have *U*^*T*^*U* = *I*_*n*_ and from the orthogonality of *V* we have *V*^*T*^*V* = *VV*^*T*^ = *I*_*n*_.

If the polarity of the *r*-th SNP is changed by swapping 0s and 1s in this *r*-th row of *Y*, this operation can be written as 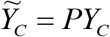, where *P* is a diagonal matrix of order *p* with *P*{*r*, *r*} = 1 if the polarity of the *r*-th SNP is unchanged and *P*{*r*, *r*} = −1 if the polarity is changed. It is important to note that *PP*^*T*^ =*P*^*T*^ *P* = *I*_*P*_. Now 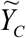 can be written as 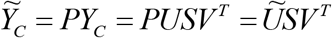, where 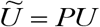. The right-hand side of this equation can be seen to represent an SVD of 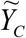 because 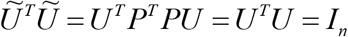. Thus, *V* is the matrix of right singular vectors of both *Y*_*C*_ and 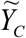. For SNP-Centered PCA, this explains why (up to a possible sign change of whole columns) the column or Individual scores remain unaltered after changing the polarity of coding (that is, swapping 0s and 1s) for any or all SNPs.

Second, consider SNP-Standardized PCA. For standardized data, *Y*_*S*_ = *D*^−1/2^*Y*_*C*_ where *D* = *diag*(*W*) with 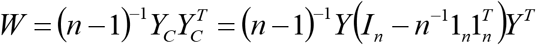. Changing the polarity of some SNPs does not change the SNP variances in *D*. Therefore, the above results for SNP-Centered data carry over fully to SNP-Standardized data.

Third and finally, consider Double-Centered PCA. DC-PCA is not immune to changes in SNP polarity as regards PCA monoplots for Individuals. Double-Centering pertains to the matrix 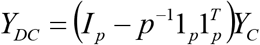. If the polarity of some SNPs are changed, then *PY*_*C*_ needs to be computed *before* the centering for Individuals. Thus, we need to compute 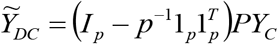. The matrices *P* and 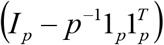 do not commute; that is, 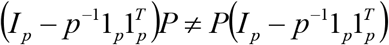 so 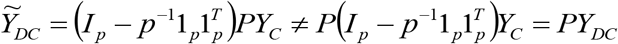. Therefore, the SVD of 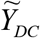 cannot be obtained from that of *Y*_*DC*_ in the same way as the SVD of 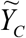 can be obtained from that of *Y*_*C*_. This explains why Individual scores before and after changing the polarity of some SNPs are not perfectly correlated.

However, when the SS for Individual main effects is small relative to that for SNP-by-Individual interaction effects, centering by Individual has little effect on the Individual scores based on SVD. The verdicts on immunity to SNP coding will be nearly the same for DC-PCA and SNP-Centered PCA when Individual main effects are small, and SNP-Centered PCA was already proven earlier in this appendix to be immune. Therefore, correlations for Individual scores between different SNP codings are expected to be very close to 1 or −1 for DC-PCA, but not exactly 1. A small SS for Individual main effects compared to that for SNP-by-Individual interaction effects is a necessary and sufficient condition for DC-PCA monoplots of Individuals to be virtually immune to changes in SNP coding.

## Acknowledgments

We appreciate helpful comments on this manuscript from Peter Bradbury, Samuel Cartinhour, Chang Chen, Kathy Esvelt Klos, Kelly Robbins, Yehao Zhang, and three anonymous reviewers. We also appreciate Kathy Esvelt Klos sharing the oat SNP data with us.

## Author Contributions

HG conceived and designed the inquiry, and invented augmented ANOVA tables. SQ analyzed data and visualized figures. SQ, LZ, and RC wrote the R software. LZ, SQ, and HG conducted the literature survey. HP wrote the appendix. HG wrote the paper with input from SQ and HP. All authors reviewed the final version of the paper.

## Supporting information

### S1 Literature Survey

Literature survey of 125 articles that apply PCA analysis to SNP data. (XLSX)

### S2 Oat Mixed

The oat dataset with SNP coding mixed as received from Kathy Esvelt Klos, except that the original coding of 1 and 2 was shifted to 0 and 1. It has 635 oat lines and 1341 SNPs. The format of this dataset is that used by our R code. (TXT)

### S3 Oat Rare1

The oat dataset with SNP coding rare=1, which required polarity reversal for 772 of the 1341 SNPs. The format of this dataset is that used by our R code. (TXT)

### S4 CA Results

CA1 arranged matrices and CA biplots for three SNP codings, using the same toy data that was used for Fig 4 in the main text. (DOCX)

### S5 Software

R code used to perform PCA and CA analyses. This R code was produced for our own in-house research purposes, not as polished and public software, but it is made available here for the sake of transparency in research. It makes basic PCA biplots and ANOVA tables, but not the polished figures and tables and the CA1 arranged matrix that appear in this publication. (R)

### S6 Four Tables

Augmented ANOVA table for SNP-Centered PCA of data on oats using SNP coding VCF with oat line 189 as the reference genome; and three augmented ANOVA tables for SNP-Standardized, Individual-Standardized, and Grand-Mean-Centered PCA using SNP coding rare=1. (DOCX)

### S7 Five Biplots

Five biplots for the oat data: SNP-Centered, SNP-Standardized, Individual-Centered, Individual-Standardized, and Grand-Mean-Centered PCA using SNP coding rare=1. As in the main text, to reduce clutter, all of these biplots use two panels, with oat lines on the left and SNPs on the right. The color scheme is the same as in Fig 1 and elsewhere in the main text, namely spring oats show in green, world diversity oats shown in blue, and winter oats shown in red, with corresponding colors for the SNPs. (DOCX)

