## Supplementary material for "Consequences of PCA graphs, SNP codings, and PCA variants for elucidating population structure": S4 CA Results

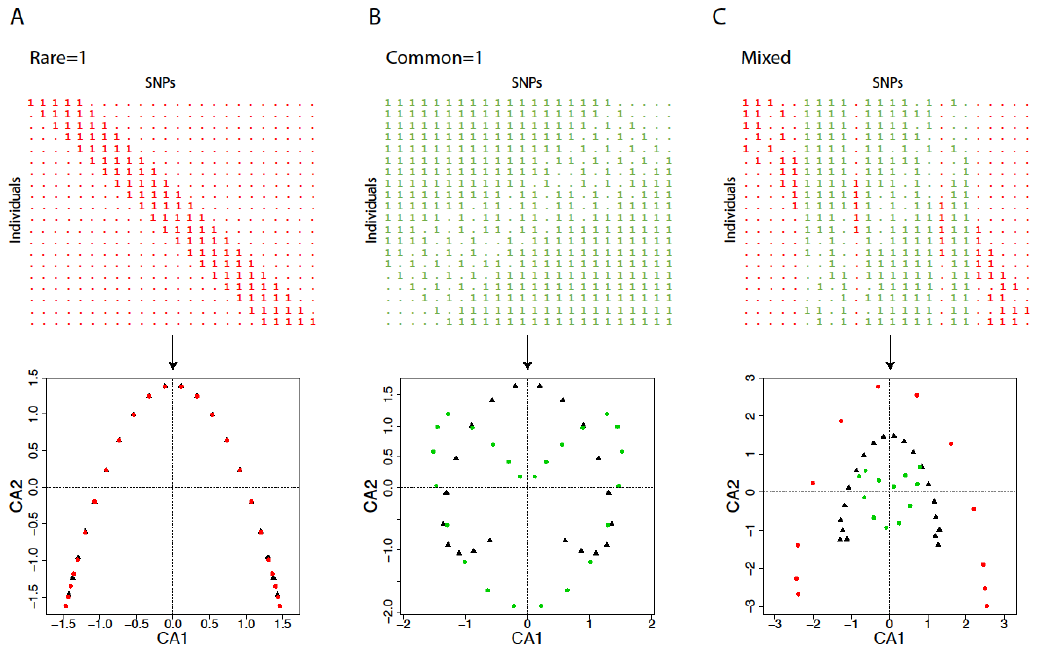

**Fig S4.1. A simple matrix and its CA biplot, using three SNP codings.** The matrix has 20 Individuals in its rows and 24 SNPs in its columns, with SNPs (columns) coded either red for SNP coding rare=1, or green for common=1. These three matrices are identical to those in Fig 4 in the main text, except that the rows and columns have been arranged in CA1 order. The biplots show the 20 Individuals as black triangles, and the 24 SNPs as dots colored either red for rare=1, or green for common=1. Unlike Fig 4, individual points are not identified because the original simple numbering does not apply to these CA1 ordered matrices. (A) For SNP coding rare=1, the CA1 ordered matrix preserves and displays the simple gradient. The CA biplot has the arch distortion, but not involuted, in contrast to the involuted arch of DC-PCA in Fig 4. (B) For SNP coding common=1, the CA1 ordered matrix does not recover the simple gradient. The CA biplot is qualitatively like the DC-PCA biplot in Fig 4 in that the arch for SNPs is rotated 180° relative to the arch for Individuals, and both arches are involuted. (C) For SNP coding mixed, the CA1 ordered matrix does not recover the simple gradient. The CA biplot has a complicated arch for the rows (black triangles) that departs at the ends of the arch from the original numbering and sequence of rows from 1 to 20 in Fig 4, and the columns (dots of either color) have a confusing pattern.
