## Supplementary material for "Consequences of PCA graphs, SNP codings, and PCA variants for elucidating population structure": S5 Software

library(ca)

#  PCA7 Six variants of PCA plus correspondence analysis (CA).  April 2019.
#  This program was written by Cornell University undergraduate students
#  Sheng Qian, Linda Zhou, and Rui Chen, supervised by Hugh G. Gauch Jr.
#
#  This software is open source and may be redistributed or modified under the
#  BSD 3-Clause License published by the Regents of the University of California.
#
#  This R code has been written for the purpose of in-house research, and it
#  has been made public for the sake of transparency.  Some researchers may find
#  this program useful, but we neither guarantee its accuracy nor provide support
#  to users.
#
#----------------------------------------------------------------------------
#  INSTRUCTIONS:
#
#  Make sure that you have installed R package "ca" in your Rstudio.
#
#  Each time you use this program, first set the working directory, ordinarily
#  to the directory that contains the datafile to be analyzed.  Also press 
#  ctrl+shift+s to run all the functions.  Then type "PCA7()" in the console 
#  below in order to run this software.
#
#----------------------------------------------------------------------------
#  DATA FILE FORMAT:
#
#  The first line has the number of columns (fastest changing subscript),
#  a name for the columns, the number of rows, a name for the rows,
#  and a name for the datase.  Data should be saved as a text file, 
#  so the example shown below could be given the filename DATA.txt.  
#  The data are numeric, with values separated by one or more spaces
#  (or a line break).
#
#  24 COLS 20 ROWS DATA
#  1 1 1 1 1 0 0 0 0 0 0 0 0 0 0 0 0 0 0 0 0 0 0 0
#  0 1 1 1 1 1 0 0 0 0 0 0 0 0 0 0 0 0 0 0 0 0 0 0
#  0 0 1 1 1 1 1 0 0 0 0 0 0 0 0 0 0 0 0 0 0 0 0 0
#  0 0 0 1 1 1 1 1 0 0 0 0 0 0 0 0 0 0 0 0 0 0 0 0
#  0 0 0 0 1 1 1 1 1 0 0 0 0 0 0 0 0 0 0 0 0 0 0 0
#  0 0 0 0 0 1 1 1 1 1 0 0 0 0 0 0 0 0 0 0 0 0 0 0
#  0 0 0 0 0 0 1 1 1 1 1 0 0 0 0 0 0 0 0 0 0 0 0 0
#  0 0 0 0 0 0 0 1 1 1 1 1 0 0 0 0 0 0 0 0 0 0 0 0
#  0 0 0 0 0 0 0 0 1 1 1 1 1 0 0 0 0 0 0 0 0 0 0 0
#  0 0 0 0 0 0 0 0 0 1 1 1 1 1 0 0 0 0 0 0 0 0 0 0
#  0 0 0 0 0 0 0 0 0 0 1 1 1 1 1 0 0 0 0 0 0 0 0 0
#  0 0 0 0 0 0 0 0 0 0 0 1 1 1 1 1 0 0 0 0 0 0 0 0
#  0 0 0 0 0 0 0 0 0 0 0 0 1 1 1 1 1 0 0 0 0 0 0 0
#  0 0 0 0 0 0 0 0 0 0 0 0 0 1 1 1 1 1 0 0 0 0 0 0
#  0 0 0 0 0 0 0 0 0 0 0 0 0 0 1 1 1 1 1 0 0 0 0 0
#  0 0 0 0 0 0 0 0 0 0 0 0 0 0 0 1 1 1 1 1 0 0 0 0
#  0 0 0 0 0 0 0 0 0 0 0 0 0 0 0 0 1 1 1 1 1 0 0 0
#  0 0 0 0 0 0 0 0 0 0 0 0 0 0 0 0 0 1 1 1 1 1 0 0
#  0 0 0 0 0 0 0 0 0 0 0 0 0 0 0 0 0 0 1 1 1 1 1 0 
#  0 0 0 0 0 0 0 0 0 0 0 0 0 0 0 0 0 0 0 1 1 1 1 1
#
#----------------------------------------------------------------------------
#
#
read <- function(name){
  file = file(name , "r")
  firstline = readLines(file, n = 1)
  # Split the string with space as an separation
  firstline = strsplit(firstline, split = " ")[[1]]
  firstline = firstline[which(firstline!="")]
  # Assertion: check if 5 items; check if intergers hold
  if(length(firstline) != 5){
    stop("Error: first line fails to have exactly five items.")
  }
  # Extract the number of the column
  NCOL = as.numeric(firstline[1])
  # Extract the name of the column
  NAME_COL = firstline[2]
  # Extract the number of the row
  NROW = as.numeric(firstline[3])
  if(NCOL%%1 != 0| NROW%%1 != 0){
    stop("First line fails to contain integer value for number of rows or columns.")
  }
  # Extract the name of the row
  NAME_ROW = firstline[4]
  # Extract the title of the dataset
  TITLE = firstline[5]
  # Read the data into a vector
  data_vector = scan(file = name, skip = 1)
  # Construct a matrix using the vector
  D = matrix(data_vector, nrow = NROW, ncol = NCOL, byrow = TRUE)
  attr = list(NCOL = NCOL, NAME_COL = NAME_COL, NROW = NROW, NAME_ROW=NAME_ROW, TITLE = TITLE)
  close(file)
  return(list(attr=attr, D=D))
}

data_transformation <- function(D,variants){
  # Assertion: stop if D is not a matrix
  stopifnot(is.matrix(D))
  # Store number of columns, number of rows
  NC = dim(D)[2];NR = dim(D)[1];NRC = NC*NR
  GM = mean(D)
  AR = rowMeans(D);AC = colMeans(D)
  SDR = apply(D,1,sd);SDC = apply(D,2,sd)
  C = D
  if(variants == 1){C = D-GM}
  if(variants == 2){C = t(t(D)-AC)}
  if(variants == 3){C = D-AR}
  if(variants == 4){C = t(t(D)-AC)-AR+GM}
  if(variants == 5){C = t(t(t(t(D)-AC))/SDC)}
  if(variants == 6){C = (D-AR)/SDR}
  if(variants == 7){C = D}
  
  # ANOVA
  # Compute sum of squares for rows and columns
  DSSR = NC*sum((AR - GM)^2);DSSC = NR*sum((AC - GM)^2);DSST = sum((D-GM)^2)
  ## Sum of squares for RxC interactions
  DSSRXC = DSST-DSSC-DSSR
  ## Degrees of freedom
  DDFT = NRC - 1;DDFR = NR - 1;DDFC = NC - 1;DDFRXC = (NR-1)*(NC-1)
  
  return(list(C = C,DSST=DSST,DSSR=DSSR,DSSC=DSSC,DDFT=DDFT,DDFR=DDFR,DDFC=DDFC, 
              DSSRXC=DSSRXC,DDFRXC=DDFRXC))
}

PCA <- function(C,variants){
  # Need call from previous function instead of re-extract from matrix C.
  NR = dim(C)[1]
  NC = dim(C)[2]
  STEST = 1e-8
  # Initialized matrix PR, PC, and vector EV.
  # Determine the limit for the number of components.
  NXL = min(NR-1, NC-1,7)
  PR = matrix(0, nrow = NR, ncol = NXL)
  PC = matrix(0, nrow =NXL, ncol = NC)
  EV = rep(0, NXL)
  # Store temporary information NX ITER STEST SFT
  temp = matrix(0, ncol = 4, nrow = NXL)
  colnames(temp) = c("NX","ITER","STEST","SFT")
  
  if(variants == 7){
    CA.P = ca(C)
    PR = CA.P$rowcoord[,1:NXL]
    PC = t(CA.P$colcoord[,1:NXL])
    EV = CA.P$sv[1:NXL]
  }
  else{
    for(NX in 1:NXL){
      SFT = 1
      ITER = 0
      # Initialize VR with uniform random function
      VR = runif(NR, min = -0.5, max = 0.5)
      SR = VR
      while(ITER<200 & SFT>STEST){
        ITER = ITER + 1
        # Compute VC from VR
        VC = t(VR)%*%C
        EVP = sum(VC^2)
        SVP = sqrt(EVP)
        VC = VC/SVP
        # Recalculate VR from VC by reverse sum
        VR = C%*%t(VC)
        EVP = sum(VR^2)
        SVP = sqrt(EVP)
        VR = VR/SVP
        SFT = max(abs(VR-SR))
        # Retain VR as SR
        SR = VR
      }
      # Store eigenvalue first
      EV[NX] = EVP
      
      SQVP = sqrt(SVP)
      VR = VR*SQVP
      VC = VC*SQVP
      
      PR[,NX] = VR
      PC[NX,] = VC
      
      # Write NX, ITER, STEST, SFT into a matrix
      temp[NX,] = c(NX, ITER, STEST, SFT)
      # Remove NX from data copy
      C = C - VR%*%VC
    }
  }
  return(list(PC = PC,PR = PR, EV = EV))
}

makeplot<-function(PC,PR,NCOL,NAME_COL,NROW,NAME_ROW,TITLE,NXL){
  #Choose the variant and PCs
  choice = as.integer(readline(prompt = "Which PCA variant do you want?  Type an integer from 1 to 7: "))
  while(!is.element(choice,c(1,2,3,4,5,6,7))){
    choice = as.integer(readline(prompt = "Please type an integer from 1 to 7: "))
  }
  target.PR = PR[,(abs(NXL*choice)-NXL+1):(NXL*abs(choice))]
  target.PC = PC[,(abs(NXL*choice)-NXL+1):(NXL*abs(choice))]
  if(sum(is.na(target.PR))==dim(target.PR)[1]*NXL & sum(is.na(target.PC))==dim(target.PC)[1]*NXL){
    print("Sorry but CA cannot be used with any negative data values. Do you want to make another graph?")
  }
  else{
    PC_choice1 = as.integer(readline(prompt = paste("Enter the PC you want for x-axis from 1 to",paste(as.character(NXL),", add a negative sign if you want to reverse the polarity: ",sep = ''))))
    while(!is.element(PC_choice1,(-NXL):NXL)){
      PC_choice1 = as.integer(readline(prompt = paste("Please type an integer from -",as.character(NXL),"to",paste(as.character(NXL),": ",sep = ''))))
    }
    PC_choice2 = as.integer(readline(prompt = paste("Enter the PC you want for y-axis from 1 to",paste(as.character(NXL),", add a negative sign if you want to reverse the polarity: ",sep = ''))))
    while(!is.element(PC_choice2,(-NXL):NXL)){
      PC_choice2 = as.integer(readline(prompt = paste("Please type an integer from -",as.character(NXL),"to",paste(as.character(NXL),": ",sep = ''))))
    }
    print("Graphing...")
    if(PC_choice1>0){
      PR1 = target.PR[,abs(PC_choice1)]
      PC1 = target.PC[,abs(PC_choice1)]
    }else if(PC_choice1<0){
      PR1 = target.PR[,abs(PC_choice1)]*(-1)
      PC1 = target.PC[,abs(PC_choice1)]*(-1)
    }
    #PC2 can also be both positive and negative
    if(PC_choice2>0){
      PR2 = target.PR[,abs(PC_choice2)]
      PC2 = target.PC[,abs(PC_choice2)]
    }else if(PC_choice2<0){
      PR2 = target.PR[,abs(PC_choice2)]*(-1)
      PC2 = target.PC[,abs(PC_choice2)]*(-1)
    }
    
    #Define the name of each choices
    if (choice==1){
      title="Grand Mean Centered"
    }else if(choice==2){
      title="Column Centered"
    }else if(choice==3){
      title="Row Centered"
    }else if(choice==4){
      title= "Double Centered (AMMI)"
    }else if(choice==5){
      title="Column Standardized" 
    }else if(choice==6){
      title= "Row Standardized" 
    }else if(choice==7) {
      title="Correspondence Analysis" 
    }
    #Make Plot
    par(mgp=c(2.5,1,0)) 
    plot(PC1,PC2,col = 'black', xlim = range(c(PC1,PR1)),ylim = range(c(PC2,PR2)),pch=17,cex.lab=1.7,cex.axis=1.3,
         xlab = paste('PC',as.character(abs(PC_choice1)),sep = ''),ylab = paste('PC',as.character(abs(PC_choice2)),sep = ''),
         main = paste(NCOL,NAME_COL,"(triangles)",NROW,NAME_ROW,"(circles)",TITLE, title ,sep = ' '),cex=0.6,asp=1)
    lines(PR1,PR2,col = 'red',type = 'p',pch=16,cex=0.6)
    abline(h = 0, v = 0, lty=2)
    print("Graph is ready, use the Export feature to save as image or pdf. Do you want to make another graph?")
    
  }
}

write_table <- function(variants,anova_result,pca_result,NCOL,NAME_COL,NROW,NAME_ROW,TITLE,NXL){
  options(scipen=20)
  data = anova_result$C
  result = pca_result
  # Grand Mean Centered
  if(variants == 1){
    SSTOTAL=rep(0,NXL);ROWSUM=rep(0,NXL);COLSUM=rep(0,NXL);INTERSUM=rep(0,NXL)
    df=rep(0,NXL)
    for(i in 1:NXL){
      C=as.matrix(result$PR[,i])%*%result$PC[i,]
      SSTOTAL[i]=sum(C^2)
      df[i]=NROW+NCOL-2*i+1
      ROWSUM[i] = sum(rowMeans(C)^2)*NCOL
      COLSUM[i] = sum(colMeans(C)^2)*NROW
      INTERSUM[i]=SSTOTAL[i]-ROWSUM[i]-COLSUM[i]
    }
    df_res= anova_result$DDFT-sum(df)
    SSTOTAL_RES=anova_result$DSST-sum(SSTOTAL)
    COLS_RES=anova_result$DSSC-sum(COLSUM)
    ROW_RES=anova_result$DSSR-sum(ROWSUM)
    INTER_RES=anova_result$DSSRXC-sum(INTERSUM)
    
    Source = c('Total')
    for(i in 1:NXL){
      chr = paste(" ",paste("PC",i,sep = ''))
      Source = c(Source,chr)
    }
    Source = c(Source,paste(" ","Residual"))
    
    df<-round(c(anova_result$DDFT,df,df_res),5)
    SS<-formatC(as.numeric(c(anova_result$DSST,SSTOTAL,SSTOTAL_RES)),digits = 5,format = 'f')
    COLS<-formatC(as.numeric(c(anova_result$DSSC,COLSUM,COLS_RES)),digits = 5,format = 'f')
    ROWS<-formatC(as.numeric(c(anova_result$DSSR,ROWSUM,ROW_RES)),digits = 5,format = 'f')
    CxR<-formatC(as.numeric(c(anova_result$DSSRXC,INTERSUM,INTER_RES)),digits = 5,format = 'f')
    table=format(data.frame(df,SS,COLS,ROWS,CxR,row.names = Source),justify="right",digits=5)
    colnames(table)<-c("df","SS",'SNPs','Individuals','SxI')
    name=paste("Augmented ANOVA for",TITLE,"analysis 1: Grand Mean Centered")
    cat(name,file= paste(TITLE,"anova6.txt",sep=""),append = TRUE,sep='\n')
    cat("-----------------------------------------------",file= paste(TITLE,"anova6.txt",sep=""),append = TRUE,sep='\n')
    cat(capture.output(table), file = paste(TITLE,"anova6.txt",sep=""),append= TRUE, sep = '\n')
    cat("-----------------------------------------------",file= paste(TITLE,"anova6.txt",sep=""),append = TRUE,sep='\n')
    cat("                                               ",file= paste(TITLE,"anova6.txt",sep=""),append = TRUE,sep='\n')
    cat("                                               ",file= paste(TITLE,"anova6.txt",sep=""),append = TRUE,sep='\n')
  }
  
  # Column Centered
  
  if(variants == 2){
    SSTOTAL=rep(0,NXL)
    ROWSUM=rep(0,NXL)
    INTERSUM=rep(0,NXL)
    df=rep(0,NXL)
    for(i in 1:NXL){
      C=as.matrix(result$PR[,i])%*%result$PC[i,]
      SSTOTAL[i]=sum(C^2)
      df[i]=NROW+NCOL-2*i
      ROWSUM[i] = sum(rowMeans(C)^2)*NCOL
      INTERSUM[i]=SSTOTAL[i]-ROWSUM[i]
    }
    df_res= anova_result$DDFT-anova_result$DDFC-sum(df)
    SSTOTAL_RES=anova_result$DSSR+anova_result$DSSRXC-sum(SSTOTAL)
    ROW_RES=anova_result$DSSR-sum(ROWSUM)
    INTER_RES=anova_result$DSSRXC-sum(INTERSUM)
    
    Source = c('Total',paste(" ",'SNP'), paste(" ","I&SxI"))
    for(i in 1:NXL){
      chr = paste("  ",paste("PC",i,sep = ''))
      Source = c(Source,chr)
    }
    Source = c(Source,paste("  ","Residual"))
    
    df<-round(c(anova_result$DDFT,anova_result$DDFC,anova_result$DDFT-anova_result$DDFC,df,df_res),5)
    SS<-formatC(as.numeric(c(anova_result$DSST,anova_result$DSSC,anova_result$DSSR+anova_result$DSSRXC,SSTOTAL,SSTOTAL_RES)),digits = 5,format = 'f')
    ROWS<-formatC(as.numeric(c("    ","    ",anova_result$DSSR,ROWSUM,ROW_RES)),digits = 5,format = 'f')
    CxR<-formatC(as.numeric(c("   ","   ",anova_result$DSSRXC,INTERSUM,INTER_RES)),digits = 5,format = 'f')
    
    table=format(data.frame(df,SS,ROWS,CxR,row.names = Source),justify="right",digits=11)
    colnames(table)<-c("df","SS",'Individuals','SxI')
    name=paste("Augmented ANOVA for",TITLE,"analysis 2: Column Centered")
    cat(name,file= paste(TITLE,"anova6.txt",sep=""),append = TRUE,sep='\n')
    cat("-----------------------------------------------",file= paste(TITLE,"anova6.txt",sep=""),append = TRUE,sep='\n')
    cat(capture.output(table), file = paste(TITLE,"anova6.txt",sep=""),append= TRUE, sep = '\n')
    cat("-----------------------------------------------",file= paste(TITLE,"anova6.txt",sep=""),append = TRUE,sep='\n')
    cat("                                               ",file= paste(TITLE,"anova6.txt",sep=""),append = TRUE,sep='\n')
    cat("                                               ",file= paste(TITLE,"anova6.txt",sep=""),append = TRUE,sep='\n')
  }
  # Row Centered
  
  if(variants == 3){
    SSTOTAL=rep(0,NXL)
    COLSUM=rep(0,NXL)
    INTERSUM=rep(0,NXL)
    df=rep(0,NXL)
    for(i in 1:NXL){
      C=as.matrix(result$PR[,i])%*%result$PC[i,]
      SSTOTAL[i]=sum(C^2)
      df[i]=NROW+NCOL-2*i
      COLSUM[i] = sum(colMeans(C)^2)*NROW
      INTERSUM[i]=SSTOTAL[i]-COLSUM[i]
    }
    df_res= anova_result$DDFT-anova_result$DDFR-sum(df)
    SSTOTAL_RES=anova_result$DSSC+anova_result$DSSRXC-sum(SSTOTAL)
    COLS_RES=anova_result$DSSC-sum(COLSUM)
    INTER_RES=anova_result$DSSRXC-sum(INTERSUM)
    
    Source = c("Total",paste(" ",'Individuals'), paste(" ","S&SxI"))
    for(i in 1:NXL){
      chr = paste("  ",paste("PC",i,sep = ''))
      Source = c(Source,chr)
    }
    Source = c(Source,paste("  ","Residual"))
    
    df<-round(c(anova_result$DDFT,anova_result$DDFR,anova_result$DDFT-anova_result$DDFR,df,df_res),5)
    SS<-formatC(as.numeric(c(anova_result$DSST,anova_result$DSSR,anova_result$DSSC+anova_result$DSSRXC,SSTOTAL,SSTOTAL_RES)),digits = 5,format = 'f')
    COLS<-formatC(as.numeric(c("    ","    ",anova_result$DSSC,COLSUM,COLS_RES)),digits = 5,format = 'f')
    CxR<-formatC(as.numeric(c("    ","    ",anova_result$DSSRXC,INTERSUM,INTER_RES)),digits = 5,format = 'f')
    
    table=format(data.frame(df,SS,COLS,CxR,row.names = Source),justify="right",digits=11)
    colnames(table)<-c("df","SS",'SNPs','SxI')
    name=paste("Augmented ANOVA for",TITLE,"analysis 3: Row Centered")
    cat(name,file= paste(TITLE,"anova6.txt",sep=""),append = TRUE,sep='\n')
    cat("-----------------------------------------------",file= paste(TITLE,"anova6.txt",sep=""),append = TRUE,sep='\n')
    cat(capture.output(table), file = paste(TITLE,"anova6.txt",sep=""),append= TRUE, sep = '\n')
    cat("-----------------------------------------------",file= paste(TITLE,"anova6.txt",sep=""),append = TRUE,sep='\n')
    cat("                                               ",file= paste(TITLE,"anova6.txt",sep=""),append = TRUE,sep='\n')
    cat("                                               ",file= paste(TITLE,"anova6.txt",sep=""),append = TRUE,sep='\n')
  }
  
  
  # Double Centered (AMMI)
  
  if(variants == 4){
    EV=rep(0,NXL)
    df=rep(0,NXL)
    for (i in 1:NXL){
      EV[i]=result$EV[i]
      df[i]=NCOL+NROW-2*i-1
    }
    EV_RES=anova_result$DSSRXC-sum(EV)
    df_res=anova_result$DDFRXC-sum(df)
    
    Source = c("Total",paste(" ",'SNPs'),paste(" ",'Individuals'),paste(" ",'SxI'))
    for(i in 1:NXL){
      chr = paste("  ",paste("IPC",i,sep = ''))
      Source = c(Source,chr)
    }
    Source = c(Source,paste("  ","Residual"))
    
    df<-round(c(anova_result$DDFT,anova_result$DDFC,anova_result$DDFR,anova_result$DDFRXC,df,df_res),5)
    SS<-formatC(as.numeric(c(anova_result$DSST,anova_result$DSSC,anova_result$DSSR,anova_result$DSSRXC,EV,EV_RES)),digits = 5,format = 'f')
    table=format(data.frame(df,SS,row.names = Source),justify="right",digits=11)
    name=paste("ANOVA for",TITLE,"analysis 4: Double Centered (AMMI)")
    cat(name,file= paste(TITLE,"anova6.txt",sep=""),append = TRUE,sep='\n')
    cat("-----------------------------------------------",file= paste(TITLE,"anova6.txt",sep=""),append = TRUE,sep='\n')
    cat(capture.output(table), file = paste(TITLE,"anova6.txt",sep=""),append= TRUE, sep = '\n')
    cat("-----------------------------------------------",file= paste(TITLE,"anova6.txt",sep=""),append = TRUE,sep='\n')
    cat("                                               ",file= paste(TITLE,"anova6.txt",sep=""),append = TRUE,sep='\n')
    cat("                                               ",file= paste(TITLE,"anova6.txt",sep=""),append = TRUE,sep='\n')
  }
  
  # Column Standardized
  
  if(variants == 5){
    SSTOTAL=rep(0,NXL)
    ROWSUM=rep(0,NXL)
    INTERSUM=rep(0,NXL)
    df=rep(0,NXL)
    SSR=0
    for(row in 1:NROW){
      SSR=SSR+sum((mean(data[row, ])^2))*NCOL
    }
    for(i in 1:NXL){
      C=as.matrix(result$PR[,i])%*%result$PC[i,]
      SSTOTAL[i]=sum(C^2)
      df[i]=NROW+NCOL-2*i
      ROWSUM[i] = sum(rowMeans(C)^2)*NCOL
      INTERSUM[i]=SSTOTAL[i]-ROWSUM[i]
    }
    df_res= anova_result$DDFT-anova_result$DDFC-sum(df)
    SSTOTAL_RES=sum(data^2)-sum(SSTOTAL)
    ROW_RES=SSR-sum(ROWSUM)
    INTER_RES=sum(data^2)-SSR-sum(INTERSUM)
    
    Source = c("I&SxI")
    for(i in 1:NXL){
      chr = paste(" ",paste("PC",i,sep = ''))
      Source = c(Source,chr)
    }
    Source = c(Source,paste(" ","Residual"))
    
    df<-round(c(anova_result$DDFT-anova_result$DDFC,df,df_res),5)
    SS<-formatC(as.numeric(c(sum(data^2),SSTOTAL,SSTOTAL_RES)),digits = 5,format='f')
    ROWS<-formatC(as.numeric(c(SSR,ROWSUM,ROW_RES)),digits = 5,format='f')
    CxR<-formatC(as.numeric(c(sum(data^2)-SSR,INTERSUM,INTER_RES)),digits = 5,format='f')
    
    table=format(data.frame(df,SS,ROWS,CxR,row.names = Source),justify="right",digits=11)
    colnames(table)<-c("df","SS",'Individuals',"SxI")
    name=paste("Augmented ANOVA for",TITLE,"analysis 5: Column Standardized")
    cat(name,file= paste(TITLE,"anova6.txt",sep=""),append = TRUE,sep='\n')
    cat("-----------------------------------------------",file= paste(TITLE,"anova6.txt",sep=""),append = TRUE,sep='\n')
    cat(capture.output(table), file = paste(TITLE,"anova6.txt",sep=""),append= TRUE, sep = '\n')
    cat("-----------------------------------------------",file= paste(TITLE,"anova6.txt",sep=""),append = TRUE,sep='\n')
    cat("                                               ",file= paste(TITLE,"anova6.txt",sep=""),append = TRUE,sep='\n')
    cat("                                               ",file= paste(TITLE,"anova6.txt",sep=""),append = TRUE,sep='\n')
  }
  
  # Row Standardized
  
  if(variants == 6){
    SSTOTAL=rep(0,NXL)
    COLSUM=rep(0,NXL)
    INTERSUM=rep(0,NXL)
    df=rep(0,NXL)
    SSC=0
    for(col in 1:NCOL){
      SSC=SSC+sum((mean(data[,col]))^2)*NROW
    }
    for(i in 1:NXL){
      C=as.matrix(result$PR[,i])%*%result$PC[i,]
      SSTOTAL[i]=sum(C^2)
      df[i]=NROW+NCOL-2*i
      COLSUM[i] = sum(colMeans(C)^2)*NROW
      INTERSUM[i]=SSTOTAL[i]-COLSUM[i]
    }
    df_res= anova_result$DDFT-anova_result$DDFR-sum(df)
    SSTOTAL_RES=sum(data^2)-sum(SSTOTAL)
    COLS_RES=SSC-sum(COLSUM)
    INTER_RES=sum(data^2)-SSC-sum(INTERSUM)
    
    Source = c("S&SxI")
    for(i in 1:NXL){
      chr = paste(" ",paste("PC",i,sep = ''))
      Source = c(Source,chr)
    }
    Source = c(Source,paste(" ","Residual"))
    
    df<-round(c(anova_result$DDFT-anova_result$DDFR,df,df_res),5)
    SS<-formatC(as.numeric(c(sum(data^2),SSTOTAL,SSTOTAL_RES)),digits = 5,format = 'f')
    COLS<-formatC(as.numeric(c(SSC,COLSUM,COLS_RES)),digits = 5,format = 'f')
    CxR<-formatC(as.numeric(c(sum(data^2)-SSC,INTERSUM,INTER_RES)),digits = 5,format = 'f')
    
    table=format(data.frame(df,SS,COLS,CxR,row.names = Source),justify="right",digits=11)
    colnames(table)<-c("df","SS",'SNPs',"SxI")
    name=paste("Augmented ANOVA for",TITLE,"analysis 6: Row Standardized")
    cat(name,file= paste(TITLE,"anova6.txt",sep=""),append = TRUE,sep='\n')
    cat("-----------------------------------------------",file= paste(TITLE,"anova6.txt",sep=""),append = TRUE,sep='\n')
    cat(capture.output(table), file = paste(TITLE,"anova6.txt",sep=""),append= TRUE, sep = '\n')
    cat("-----------------------------------------------",file= paste(TITLE,"anova6.txt",sep=""),append = TRUE,sep='\n')
    cat("                                               ",file= paste(TITLE,"anova6.txt",sep=""),append = TRUE,sep='\n')
    cat("                                               ",file= paste(TITLE,"anova6.txt",sep=""),append = TRUE,sep='\n')
  }
}

plot_request<-function(flag,PC,PR,NCOL,NAME_COL,NROW,NAME_ROW,TITLE,NXL){
  want_graph=readline(prompt="Want graphs? Type y/n:")
  while(flag==TRUE){
    if(want_graph =="y"|want_graph=="Y"){
      makeplot(PC,PR,NCOL,NAME_COL,NROW,NAME_ROW,TITLE,NXL)
      want_graph=readline(prompt="Type y/n:")
    }
    else if (want_graph == "n"|want_graph=="N"){
      print("Thanks for using PCA software!")
      flag=FALSE
    }
    else{
      print("Invalid command")
      plot_request(flag,PC,PR,NCOL,NAME_COL,NROW,NAME_ROW,TITLE,NXL)
      flag=FALSE
    }
  }
}

PCA7<-function(){
  set_directory=readline(prompt="Have you set your working director? Type y/n:")
  if(set_directory =="y"|set_directory=="Y"){
    print("Choose the data file in your browser.")
    the_file = read(file.choose())
    NXL = min(dim(the_file[[2]])[1]-1, dim(the_file[[2]])[2]-1,7)
    PR = matrix(NA,nrow=dim(the_file[[2]])[1],ncol = 7*NXL)
    PC = matrix(NA,nrow=dim(the_file[[2]])[2],ncol = 7*NXL)
    print("Read data. Analyses begin.")
    NCOL= the_file$attr[[1]]
    NAME_COL= the_file$attr[[2]]
    NROW=the_file$attr[[3]]
    NAME_ROW = the_file$attr[[4]]
    TITLE=the_file$attr[[5]]
    # MAIN FUNCTION STARTS HERE
    for(i in 1:7){
      anova_result = data_transformation(the_file[[2]],i)
      data = anova_result$C
      if (i==7){
        if (sum(data<0)!=0){
          print("Correspondence analysis is not available because some data are negative.")
          next
        }
      }
      pca_result = PCA(data,i)
      pr = pca_result$PR
      pc = pca_result$PC
      low = (NXL*i-NXL+1)
      upper = (NXL*i)
      if(is.null(dim(pr))){
        PR[,low:upper]= pr
      }
      else{
        PR[,low:upper]= pr[,1:NXL]
      }
      if(is.null(dim(pc))){
        PC[,low:upper]= pc
      }
      else{
        PC[,low:upper]= t(pc[1:NXL,])
      }
      write_table(i,anova_result,pca_result,NCOL,NAME_COL,NROW,NAME_ROW,TITLE,NXL)
      if (i==1){
        title="Grand Mean Centered"
      }else if(i==2){
        title=paste("Column"," Centered",sep="")
      }else if(i==3){
        title=paste("Row"," Centered",sep="")
      }else if(i==4){
        title= "Double Centered (AMMI)"
      }else if(i==5){
        title=paste("Column"," Standardized",sep="")
      }else if(i==6){
        title= paste("Row"," Standardized",sep="")
      }else if(i==7) {
        title="Correspondence Analysis" 
      }
      print(paste("Finished analysis",i,"of 7:",title))
    }
    print("Finished all 7 analyses.")
    print("Tables are ready in the working directory.")
    
    # Make Graphs
    flag=TRUE
    plot_request(flag,PC,PR,NCOL,NAME_COL,NROW,NAME_ROW,TITLE,NXL)
  }
  else if (set_directory == "n"|set_directory=="N"){
    print("Please set your working directory!")
    flag=FALSE
  }
  else{
    print("Invalid command")
  }
}
