## Supplementary material for "Consequences of PCA graphs, SNP codings, and PCA variants for elucidating population structure": S6 Four Tables

**Table S6.1. Augmented ANOVA table for SNP-Centered PCA of oat data, using SNP coding VCF with oat line 189 as the reference genome.**

```
———————————————————————————————————————————————————————————
Source            df           SS  Individuals          SxI
———————————————————————————————————————————————————————————
Total         851534   175776.962
  SNPs          1340    35206.937
  I&SxI       850194   140570.025     6286.978   134283.047
    PC1         1974    16325.612     5072.449    11253.163
    PC2         1972     9751.158      279.884     9471.275
    PC3         1970     6250.504       78.151     6172.353
    PC4         1968     4860.910      503.165     4357.745
    PC5         1966     3510.312       16.352     3493.959
    PC6         1964     3181.953       45.154     3136.799
    PC7         1962     2891.656       23.786     2867.870
    Residual  836418    93797.920      268.037    93529.883
———————————————————————————————————————————————————————————
```

**Table S6.2. Augmented ANOVA table for SNP-Standardized PCA of oat data, using SNP coding rare=1.**

```
————————————————————————————————————————————————————————
Source          df          SS  Individuals          SxI
————————————————————————————————————————————————————————
I&SxI       850194  850194.000    14427.785   835766.215
  PC1         1974   84215.540     5870.564    78344.976
  PC2         1972   54117.580     1751.478    52366.102
  PC3         1970   34779.168     3713.961    31065.207
  PC4         1968   29068.093      279.528    28788.565
  PC5         1966   23181.728      168.888    23012.840
  PC6         1964   21843.243        0.403    21842.840
  PC7         1962   17066.966      591.352    16475.614
  Residual  836418  585921.681     2051.609   583870.072
————————————————————————————————————————————————————————
```

**Table S6.3. Augmented ANOVA table for Individual-Standardized PCA of oat data, using SNP coding rare=1.**

```
————————————————————————————————————————————————————————
Source          df          SS         SNPs          SxI
————————————————————————————————————————————————————————
S&SxI       850900  850900.000    93428.621   757471.379
  PC1         1974   98971.642    81776.097    17195.545
  PC2         1972   79236.782    11285.246    67951.536
  PC3         1970   49788.726        2.118    49786.608
  PC4         1968   33060.462        1.813    33058.649
  PC5         1966   27932.581      215.481    27717.100
  PC6         1964   19288.835       25.445    19263.390
  PC7         1962   16246.322        5.008    16241.314
  Residual  837124  526374.649      117.412   526257.237
————————————————————————————————————————————————————————
```

**Table S6.4. Augmented ANOVA table for Grand-Mean-Centered PCA of oat data, using SNP coding rare=1.**

```
—————————————————————————————————————————————————————————————————————
Source          df          SS  Individuals         SNPs          SxI
—————————————————————————————————————————————————————————————————————
Total       851534  157442.756     2077.808    16872.731   138492.217
  PC1         1975   17402.677        0.490    16741.733      660.454
  PC2         1973   16325.068      859.333        7.902    15457.833
  PC3         1971    9709.637      252.328       48.779     9408.530
  PC4         1969    6217.325      441.158       16.201     5759.966
  PC5         1967    4774.109       12.132       29.935     4732.042
  PC6         1965    3490.992       11.347        4.086     3475.559
  PC7         1963    3144.600      174.890        7.697     2962.013
  Residual  837751   96378.348      326.130       16.398    96035.820
—————————————————————————————————————————————————————————————————————
```
