## Supplementary material for "Consequences of PCA graphs, SNP codings, and PCA variants for elucidating population structure": S7 Five Biplots

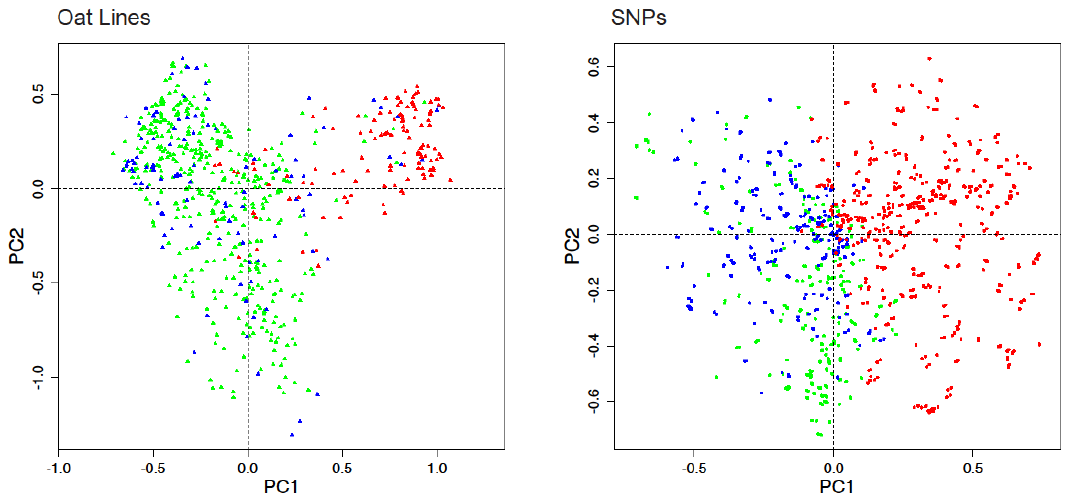
These two figures show biplots for SNP-Centered and SNP-Standardized PCA for the oat data, using SNP coding rare=1. They are quite similar, and are nearly identical for both panels to Fig 1 in the main text, which uses DC-PCA and the same SNP coding.

**Fig S7.1. SNP-Centered PCA for the oat data, using SNP coding rare=1.**

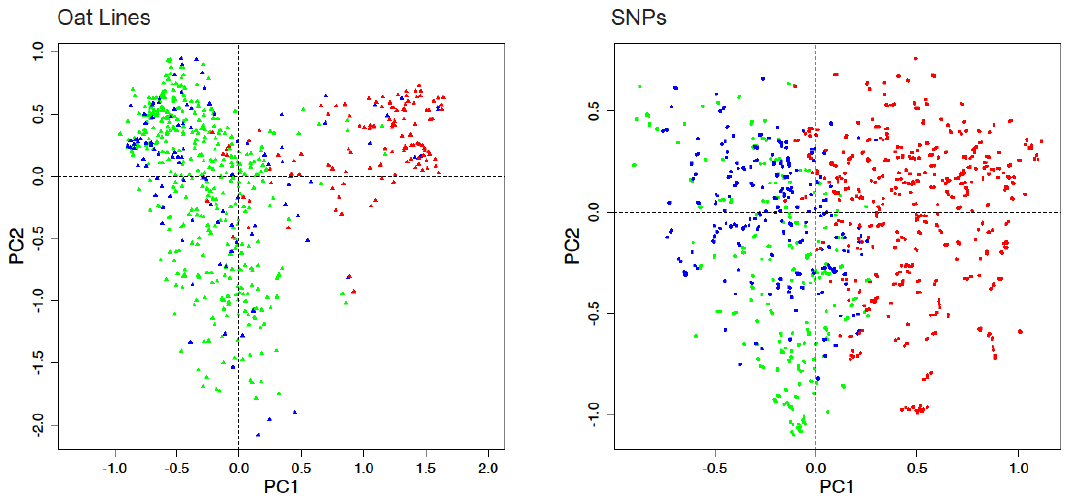

**Fig S7.2. SNP-Standardized PCA for the oat data, using SNP coding rare=1.**

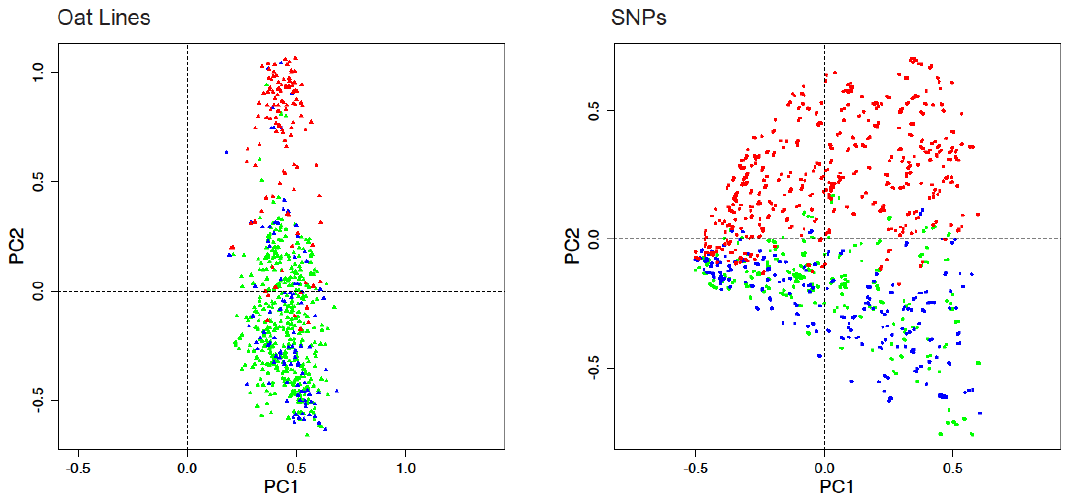
These two figures show biplots for Item-Centered and Item-Standardized PCA for the oat data, using SNP coding rare=1. They are similar, and are also similar for both panels to Fig 7 in the main text, which uses AMMI1 and the same SNP coding.

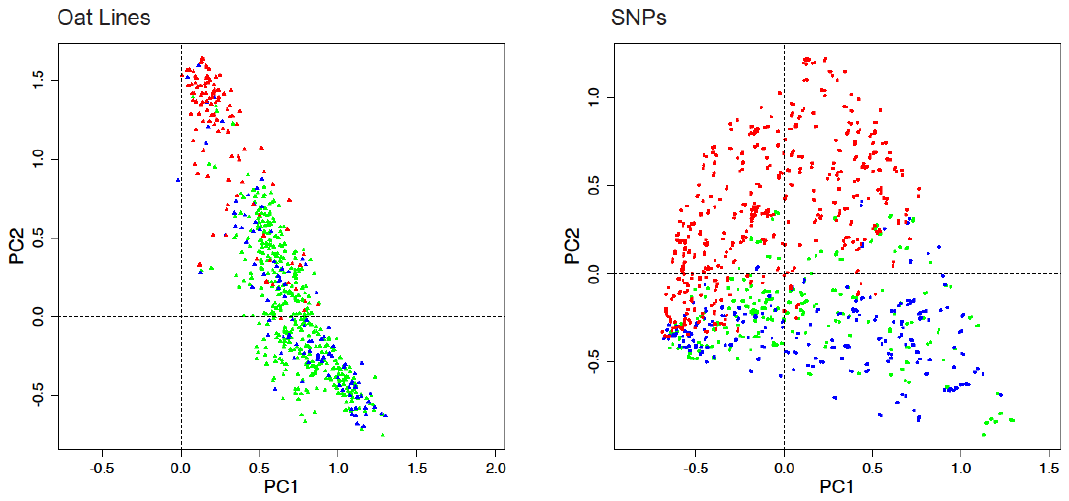
**Fig S7.3. Item-Centered PCA for the oat data, using SNP coding rare=1.**

**Fig S7.4. Item-Standardized PCA for the oat data, using SNP coding rare=1.**

This figure shows a biplot for Grand-Mean-Centered PCA for the oat data, using SNP coding rare=1. This biplot is similar for both panels to Fig 7 in the main text, which uses AMMI1 and
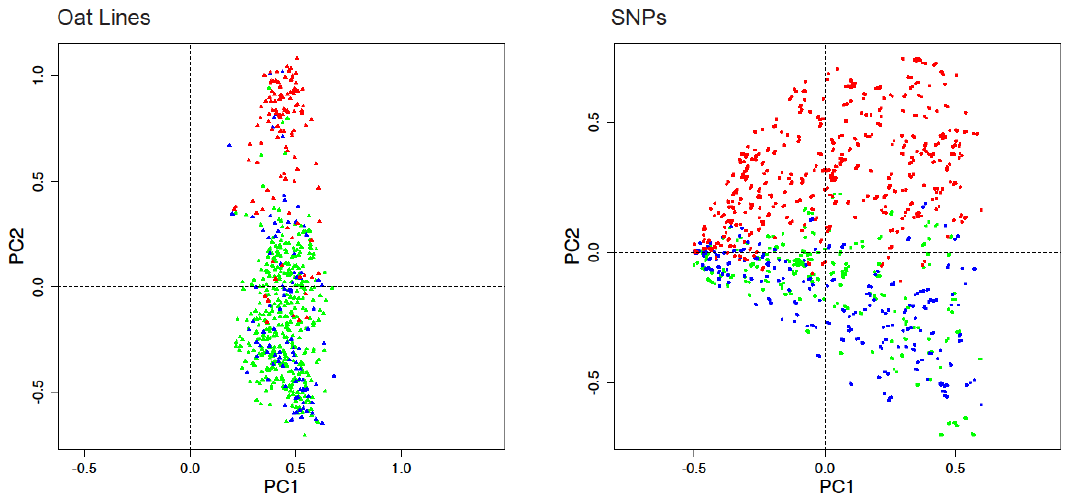
the same SNP coding.

**Fig S7.5. Grand-Mean-Centered PCA for the oat data, using SNP coding rare=1.**
